# Single-cell transcriptomics reveals cellular hierarchies and aberrant CTS contraction-mediated premature hair regression in androgenetic alopecia

**DOI:** 10.1101/2023.12.25.573294

**Authors:** Guo Li, Li Yang, Shixin Duan, Mengting Chen, Yujin Zhang, Fangfen Liu, Yan Tang, Yunying Wang, Jiayun Li, San Xu, Zheng Wu, Ben Wang, Zhixiang Zhao, Wei Shi, Hongfu Xie, Zhili Deng, Ji Li

## Abstract

Androgenetic alopecia (AGA) is characterized by progressive miniaturization of hair, forming a distinctive patterned baldness in the scalp; yet, the mechanisms for hair miniaturization in this disease remain largely unknown. Here by single-cell transcriptome analysis, we describe a concise single-cell atlas, and identify the early changes in cell subpopulations, hair follicle (HF) stem cell fate determination and cell-cell communications in AGA anagen HF units. Thereinto, apoptotic loss of HF progenitor cells is significantly increased, correlated with HF miniaturization in AGA. Mechanistically, enhanced contraction of connective tissue sheath (CTS) activates the mechanosensitive channel PIEZO1, which triggers ectopic apoptosis of progenitor cells in human anagen HFs. Continuous CTS contraction during AGA causes long-term loss of progenitor cells via inducing persistent ectopic apoptosis through PIEZO1, eventually leading to premature hair regression. Most importantly, we show that targeting CTS contraction by ML-7, a selective myosin light chain kinase (MLCK) inhibitor, can obviously improve the growth of HFs from balding scalps of AGA patients. Our study reveals the cellular hierarchies and identifies CTS with increased muscle contraction activity as a driver of premature hair regression in AGA, highlighting CTS surrounding human HF as the therapeutic target for treating this disorder.

## Introduction

Androgenetic alopecia (AGA), also known as male pattern baldness, is the most common type of baldness, affecting both men and women. Testosterone and androgen receptor (AR) signaling pathway are considered to be necessary for AGA development(*1, 2*). Genetic susceptibility loci mapping to AR gene and its neighboring ectodysplasin A2 receptor gene (EDA2R) have also been found in AGA(*3*). Current recognized therapies for AGA include finasteride, minoxidil, platelet-rich plasma (PRP) and low-level laser light therapy (LLLT)(*4, 5*). Among these, finasteride suppresses the activity of 5-a reductase 2 (SRD5A2), which transforms testosterone to dihydrotestosterone (DHT), a more potent AR agonist(*1*). Although the potential targets of other therapies remain elusive, additional factors are suggested to be involved in the development of AGA besides AR-related pathways; however, specific instances have not been conclusively identified.

AGA is featured by the progressive and patterned conversion of large terminal hair follicles (HFs) into small vellus HFs (termed as HF miniaturization), which eventually results in balding(*6*). All HFs undergo periodic cycles of growth (anagen), regression (catagen), and rest (telogen) stages throughout the lifetime(*7*). Normally, most human hair follicles will remain in anagen for 2-6 years to produce large terminal hair. But in AGA, the growth phase decreases to only days or weeks, followed by premature entry into hair regression; this leads to an increase in the percentage of resting HFs, generating microscopic hairs in the balding scalp as the HF cycles(*8, 9*). In HF cycle, the growth of HFs relies on the proliferation and differentiation of HF stem/progenitor cells, whose activities are mainly regulated by the components of HF niche (including dermal papilla (DP), dermal sheath (DS), adipocytes and blood vessels *etc*.)(*6, 10–13*). In balding scalp of AGA, HF stem cells (HFSCs), also known as bulge cells, are basically retained, whereas the number of progenitor cells, a cluster of more actively proliferating cells, is decreased(*14*). This suggests that HFs in AGA balding scalp lack fuels for growth, but the details and underlying mechanism are unclear.

To date, the pathogenetic study of AGA has been scant and limited to histomorphology, global gene analysis and immunohistochemistry due to the fact that most laboratory models are not fully representive of AGA(*15, 16*). Single-cell RNA-sequencing (scRNA-seq) has emerged as a powerful tool for dissecting the transcriptome dynamics, cell heterogeneity, cell lineage tracing and cell-cell communications in complex tissues at uniquely high resolution(*17–21*). However, to our knowledge, a study regarding scRNA-seq application for exploring the pathogenesis of AGA has never been described.

In this study, we report a comprehensive scRNA-seq analysis on human anagen HF units from 3 conditions: balding frontal and non-balding occipital scalps of patients with AGA, and normal frontal scalps of healthy individuals. With these data, we generated a concise single-cell atlas and elucidated the early cellular and molecular changes in AGA anagen HF units. Importantly, via combining human HF organ culture system, we revealed an essential role of connective tissue sheath (CTS), the analogous structure of murine DS in human, in regulating HF progenitor cell survival in the development of AGA. In AGA, CTS contraction is significantly enhanced, resulting in mechanical compression of HF progenitor cells; increased mechanical force activates the mechanosensitive calcium channel PIEZO1, thus induces apoptotic loss of progenitor cells, which eventually leads to premature hair regression. Finally, we demonstrated that targeting CTS contraction by ML-7 can promote the growth of HFs from the balding scalps of AGA patients, even better than minoxidil. These findings illustrate the cellular hierarchies in AGA and emphasize relaxing hair follicle by targeting CTS as a promising strategy for its treatment.

## Results

### Single-cell RNA-seq reveals cell heterogeneity of androgenetic alopecia and healthy anagen HF units

To dissect the cellular heterogeneity and explore the early changes in AGA development, we obtained anagen HFs from frontal balding (B) scalps and occipital non-balding (NB) scalps of AGA patients, and frontal normal scalps of healthy individuals (HS), to perform scRNA-seq (Fig. 1A). Among these, anagen HFs of balding scalps were at the early stage of hair miniaturization, isolated from the edge of frontal hairline of AGA patients. First, to identify the hair cycle phase of human HFs, we established an identification strategy by integrating multiple methods reported previously(*22, 23*) (Fig. S1). Before conducting scRNA-seq, the HFs were confirmed in anagen phase. After sequencing and stringent cell filtration, 76368 cells were retained for subsequent analyses. Following a standard scRNA-seq data analysis procedure, totally 28 clusters (C0-C27) were retained, which were sufficiently separated from each other (Fig. S2A), and these clusters could be reproduced using cells from each of the 12 samples, suggesting that they are robustly present across different samples and conditions (including HS, NB and B) (Fig. S2B, C).

**Fig. 1:**
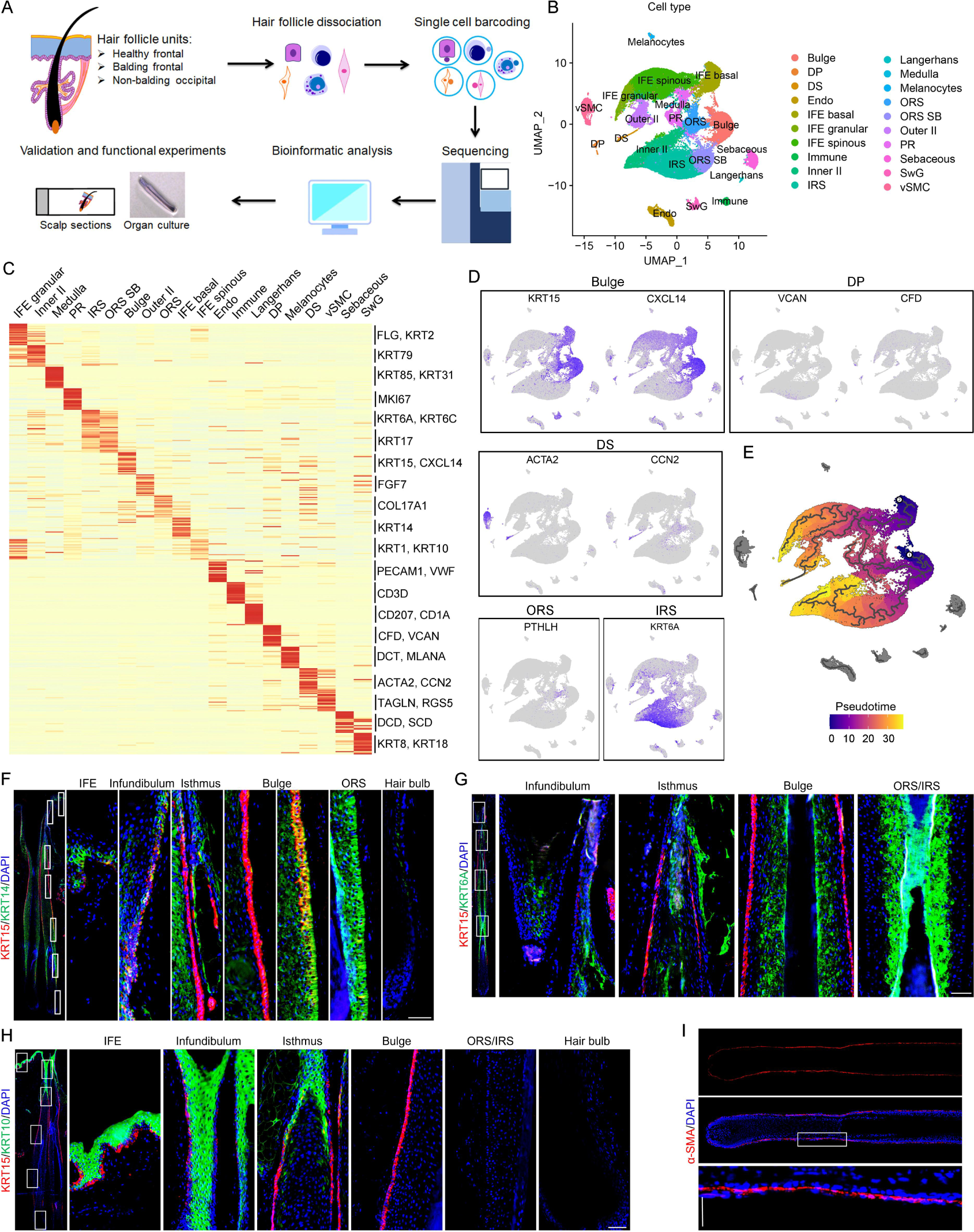
scRNA-seq reveals cell type composition in anagen HF units of AGA and healthy scalps. **A** Flowchart overview of scRNA-seq, and subsequent validation and functional experiments with anagen HFs from balding frontal and non-balding occipital scalps of AGA patients, and normal scalps of healthy individuals. **B** Uniform manifold approximation and projection (UMAP) plot showing the twenty cell types of human anagen HFs. DP, dermal papilla; DS, dermal sheath; Endo, endothelial cells; IFE, interfollicular epidermis; Inner II, inner layer of infundibulum and isthmus; IRS, inner root sheath; ORS, outer root sheath basal; ORS SB, outer root sheath suprabasal; Outer II, outer layer of infundibulum and isthmus; PR, proliferating progenitors; SwG, sweat gland; vSMC, vascular smooth muscle cells. **C** Heatmap of top 20 marker genes of each HF cell type. **D** Feature plots showing expression of marker genes for the Bulge (KRT15, KRT14), DP (VCAN, CFD), DS (ACTA2, CCN2), ORS (PTHLH), IRS (KRT6A). **E** Monocle3 pseudotime of all HF cell types. **F-I** Representative immunostaining images of KRT15/KRT14 (**F**), KRT6A/KRT15 (**G**), KRT15/KRT10 (**H**) and α-SMA (**I**) in human scalp sections and the corresponding magnified views of boxed areas. (n=5 HFs from 3 individuals. The same 3 individuals’ samples were used in all following stainings unless otherwise noted). Scale bar: 50 μm.

To dissect the cell types, we first defined the expression pattern of SOX9 (Fig. S2D), a reported marker for hair follicle cells(*24*). By immunohistochemistry, we confirmed that SOX9 is expressed in most cells of hair follicle and almost none in other cells, distinguishing hair follicle cells from other epidermal and dermal cells (Fig. S2E). Based on the expression of SOX9, the above clusters were further identified as 20 different cell types, according to the expression of known lineage markers, including Bulge (KRT15^+^, CXCL14^+^), DP (VCAN^+^, CFD^+^), DS (ACTA2^+^, CCN2^+^), outer root sheat basal (ORS, KRT14^+^, PTHLH^+^), ORS suprabasal (ORS SB, KRT17^+^), inner root sheat (KRT6A^+^), medulla (KRT85^+^, KRT31^+^), proliferating progenitors (PR, MKI67^+^), endothelial cells (Endo, PECAM1^+^, VWF^+^), immune cells (CD3D^+^), Langerhans (CD207^+^, CD1A^+^), Melanocytes (DCT^+^, MLANA^+^), Vascular smooth muscle cells/pericytes (vSMC, ACTA2^+^, RGS5^+^, RERGL^+^), Sebaceous gland cells (DCD^+^, SCD^+^), sweet gland cells (SwG, KRT8^+^, KRT18^+^), interfollicular epidermis spinous (IFE spinous, KRT1^+^, KRT10^+^), IFE granular (FLG^+^, KRT2^+^); cells highly expressing KRT14 but not SOX9 were defined as IFE basal; Inner II (KRT79^+^) is the outer layer of Infundibulum and Isthmus which highly expressed KRT14 and KRT6A in the upper HF region; in contrast, Outer II has a relatively low expression of KRT6A(*25–35*) (Fig. 1B-D; Supplementary Table 1; Fig. S2F-K).

To investigate the trajectories among all 20 cell types, monocle3(*36*) was employed to perform the pseudotime analysis with the root node setting at start point of bulge on the SOX9^+^ HF branch and start point of IFE basal on the SOX9^-^ IFE branch separately. Our data demonstrate that bulge cells first differentiate into ORS and IRS cells, which then further branch into SOX9^+^ PR, also known as matrix cells, and finally contribute to medulla (HF) formation. For the SOX9^-^ IFE branch, IFE basal cells can directly differentiate into SOX9^-^ PR and IFE spinous cells which finally develop into IFE granular cells (Fig. 1E). These patterns are consistent with the knowledge from previous studies(*26, 35*). Consistent with the scRNA-seq data, immunostaining analysis showed that KRT15 was highly expressed in bulge HFSCs and moderately expressed in IFE basal cells (Fig.1F), similar to previous observations(*14, 26*). As an IRS marker, KRT6A was exactly expressed mainly in the IRS and some of the inner layer cells of the infundibulum and isthmus, which are the upward continuation of IRS (Fig.1G and Fig. S2L). Moreover, we verified IFE spinous cells with KRT10 and DS cells with α-SMA (Fig. 1H, I). We determined the relative proportion of each cell type in all samples, showing increased DS and sebaceous gland cells in B anagen HF units compared with NB or HS (Fig. S2M). We next explored the number of differentially expressed genes (DEGs) between B and NB/HS. The results showed that all cell types had differences in various degree (Fig. S2N, Supplementary Table 2), suggesting that anagen HFs undergo significant changes at the early stage of hair miniaturization in AGA.

To further explore the potential heterogeneity, bulge cells were subclustered into three distinct subclusters (Fig. 2A), in which the percentage of subcluster 1 was reduced in the balding bulge of AGA patients (Fig. 2B). Wilcoxon rank-sum test was used to characterize the marker genes specifically enriched in each subpopulation versus other subpopulations. Top marker genes with avg_log2FC>0.25, pct (minimum fraction in either of the two populations)>0.25 and P value<0.05 of each subcluster were showed in Fig. S3A and Supplementary Table 3. Among these, KRT15 was commonly highly expressed (Fig. 2C); CD200 was visibly enriched in subcluster 0 (Fig. 2D), while CD34 was mainly enriched in subcluster 1 (Fig. 2E). Consistently, CD34^+^ bulge cells were decreased in the bulge of balding anagen HFs (Fig. 2F). By immunostaining, we defined that CD200^+^ cells are located in the upper bulge with high stemness (known as HFSCs), while CD34 is mainly expressed in the lower bulge cells referred to as HF progenitor cells, which are differentiated from the upper bulge HFSCs (Fig. 2G, H). Via RNA Velocyto analysis, we further confirmed that CD200^+^ bulge cells are the HFSCs, which have the ability to differentiate into CD34^+^ progenitor cells (Fig. 2I).

**Fig. 2:**
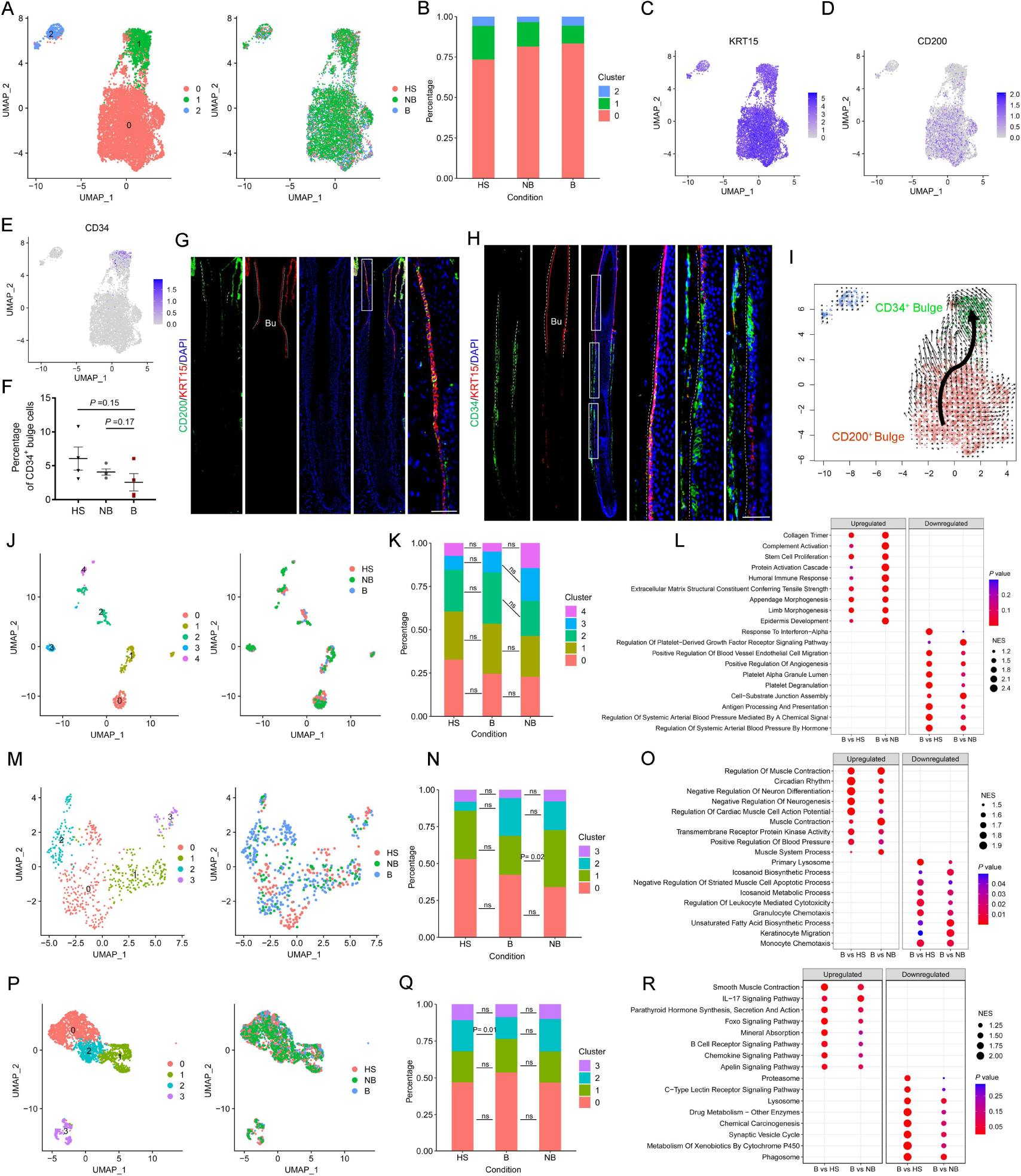
Heterogeneity of cell subpopulations in AGA and healthy scalps. **A** UMAP plots showing subclusters and sample conditions of bulge cells. **B** Bar graph representing the percentage of subpopulations of bulge cells. **C-E** Feature plots showing the expression of the KRT15 (**C**), CD200 (**D**) and CD34 (**E**) in bulge cells. **F** The ratio of CD34^+^ bulge cells in Healthy (HS), non-balding (NB) and balding (B) conditions. **G, H** Representative images showing CD200 protein expression in upper bulge stem cells (**G**) and CD34 in progenitor cells (**H**). Right panels, magnified images of boxed areas. Bu, bulge. Red, KRT15 (K15) staining. DAPI staining (blue) indicates nuclear localization. ORS–DS boundary is indicated by dashed lines. Scale bar: 50 μm. **I** velocyto showing the single-cell trajectories of bulge subclusters in UMAP. **J** UMAP plots showing subclusters and sample conditions of DP cells. **K** Bar graph representing the percentage of subpopulations of DP cells. **L** Representative GO terms of upregulated (left) and downregulated (right) DEGs between balding and healthy groups (B vs HS), balding and non-balding groups (B vs NB) in DP cells. NES stands for enrichment scores. The color keys from red to blue indicate the range of P value. **M** UMAP plot showing subclusters and sample conditions of DS cells. **N** Bar graph representing the percentage of subpopulations of DS cells. **O** Representative GO terms enriched in balding versus non-balding or healthy DS cells. NES stands for enrichment scores. The color keys from red to blue indicate the range of p value. e. **P** UMAP plot showing subclusters and sample conditions of vSMC cells. **Q** Bar graph representing the percentage of subpopulations of vSMC cells. **R** Representative GO terms enriched in balding versus non-balding or healthy vSMC cells. NES stands for enrichment scores. The color keys from red to blue indicate the range of p value. The data represent the means±SEM. P values were determined by unpaired (B vs HS) or paired (B vs NB) student’s t-test. ns, not significant.

PR cells are the proliferating progenitors which highly expressed MKI67. Further subclustering analysis separated PR cells into 2 subclusters (Fig. S3B). Cell composition analysis showed the subcluster 1 decreased in B anagen HFs of AGA (Fig. S3C). Among these two subclusters, one was enriched for epithelial markers (KRT1 and KRT10), representing the IFE proliferating progenitor (ePR) cells; another one (subcluster 1) was specifically enriched for HF cell marker SOX9, suggesting that it represented the HF proliferating progenitors also named as matrix cells (Fig. S3D, E, and Supplementary Table 4).

Subclustering of DP cells, the signaling center for hair growth(*11*), revealed 5 subclusters (Fig. 2J, Fig. S3F and Supplementary Table 5). The percentage and distribution of subclusters were comparable across HS, NB and B, suggesting that DP cell identity may not alter in AGA anagen HFs (Fig. 2K and Supplementary Table 6). Gene ontology (GO) analysis showed that angiogenesis-related pathway was downregulated (Fig. 2L), consistent with our previous findings from bulk RNA sequencing(*6*).

DS cells were divided into 4 subclusters (Fig. 2M, Fig. S3G and Supplementary Table 7), which showed no significant changes in the balding anagen HFs of AGA patients (Fig. 2N and Supplementary Table 8). Pathway enrichment analysis of differential genes revealed muscle contraction was obviously upregulated in balding anagen HFs compared to non-balding or healthy groups (Fig. 2O).

In human hair follicle, the DS is tightly surrounded by a network of blood vessels, composed of vSMCs and endothelial cells(*37*). Among them, vSMCs were separated into 4 subclusters, and the percentage and distribution of subclusters were comparable across three groups (Fig. 2P, Q; Fig. S3H; Supplementary Table 9 and Supplementary Table 10). Pathway enrichment analysis showed that smooth muscle contraction and multiple pathways involved in inflammatory regulation (such as IL-17 signaling pathway) were upregulated in vSMCs of balding anagen HFs (Fig. 2R). Endothelial cells were divided into 4 subclusters, which exhibited no obvious alterations (Fig. S3I-K; Supplementary Table 11 and Supplementary Table 12), and IL-17 signaling pathway was also upregulated in endothelial cells of balding anagen HFs (Fig. S3L).

Immune cells expressing CD3D were further separated into 6 subclusters, including Th17 cells (IL17A^+^, IL17F^+^ and IL22^+^), naive CD4^+^ T cells (IL7R^+^), CD8^+^ T cells (GZMK^+^ and CD8A^+^), natural killer T (NKT) cells (ZNF683^+^) and two unknown subclusters (Fig. S3M-O; Supplementary Table 13). Cell composition analysis showed significant increase of Th17 cells in balding HF units (Fig. S3P, Q), which provides evidence for inflammatory infiltration in AGA.

### Identification of early changes in HFSC fate determination and cell-cell communications in AGA

Epidermal and sebaceous differentiation of HFSCs has been reported to be associated with HF miniaturization during ageing-induced and obesity-induced hair loss in mice(*38, 39*), but it was unclear whether there existed HFSC fate determination alterations in AGA development. To address this question, we conducted cell fate analysis among potential cell types: bulge, ORS, sebaceous gland and IFE basal cells. As expected, more HFSCs showed a pronounced tendency to differentiate into sebaceous gland and IFE basal cells in balding HFs compared with non-balding and healthy HFs, and obviously decreased HFSCs differentiated into ORS cells (Fig. S4A, D). To explore the underlying mechanisms, branched expression analysis modeling (BAEM) was performed, and identified three common cell fate determination genes, COL17A1, PPP1R1C and SGK1 (Fig. S4B, C, E, F).

To further characterize the cell-cell communications between different cell types in HF units. Cellchat(*21*) was employed to compare the cell-cell communications between balding and non-balding/healthy HFs. Although the interaction strength showed that most cell types had no obvious changes in incoming and outgoing interaction strength, Th17 cells and langerhans exhibited increased incoming interaction strength in balding and non-balding HFs compared to healthy HFs, while ORS cells displayed decreased outgoing interaction strength in balding anagen HFs in AGA (Fig. S4G).

### Apoptotic loss of HF progenitor cells is correlated with HF miniaturization in AGA

AGA is characterized by the progressive conversion of the terminal (t) scalp HFs into intermediate/miniaturized (i/m) HFs, which we called the process of HF miniaturization and may be attributed to the deficiency of progenitor cells based on our scRNA-seq data as mentioned above and the early study(*14*). To verify changes in HFSC and progenitor cell numbers, we performed immunostaining of CD200 and CD34 on anagen HFs from frontal balding (B) and occipital non-balding scalps (NB) of AGA patients, and frontal normal scalps of healthy individuals (HS). HFs retain CD200^+^ HFSCs with the same percentage in AGA, despite a decrease in cell numbers in B scalps (Fig. S5A-C). However, CD34^+^ progenitor cells were gradually diminished during the process of HF miniaturization (Fig. 3A-C). Further analysis indicated a strong positive correlation between the number/ proportion of CD34^+^ progenitor cells and the hair follicle diameter in balding scalps of AGA patients (Fig. 3D-G). Collectively, these observations suggest that loss of progenitor cells is associated with HF miniaturization in AGA.

**Fig. 3:**
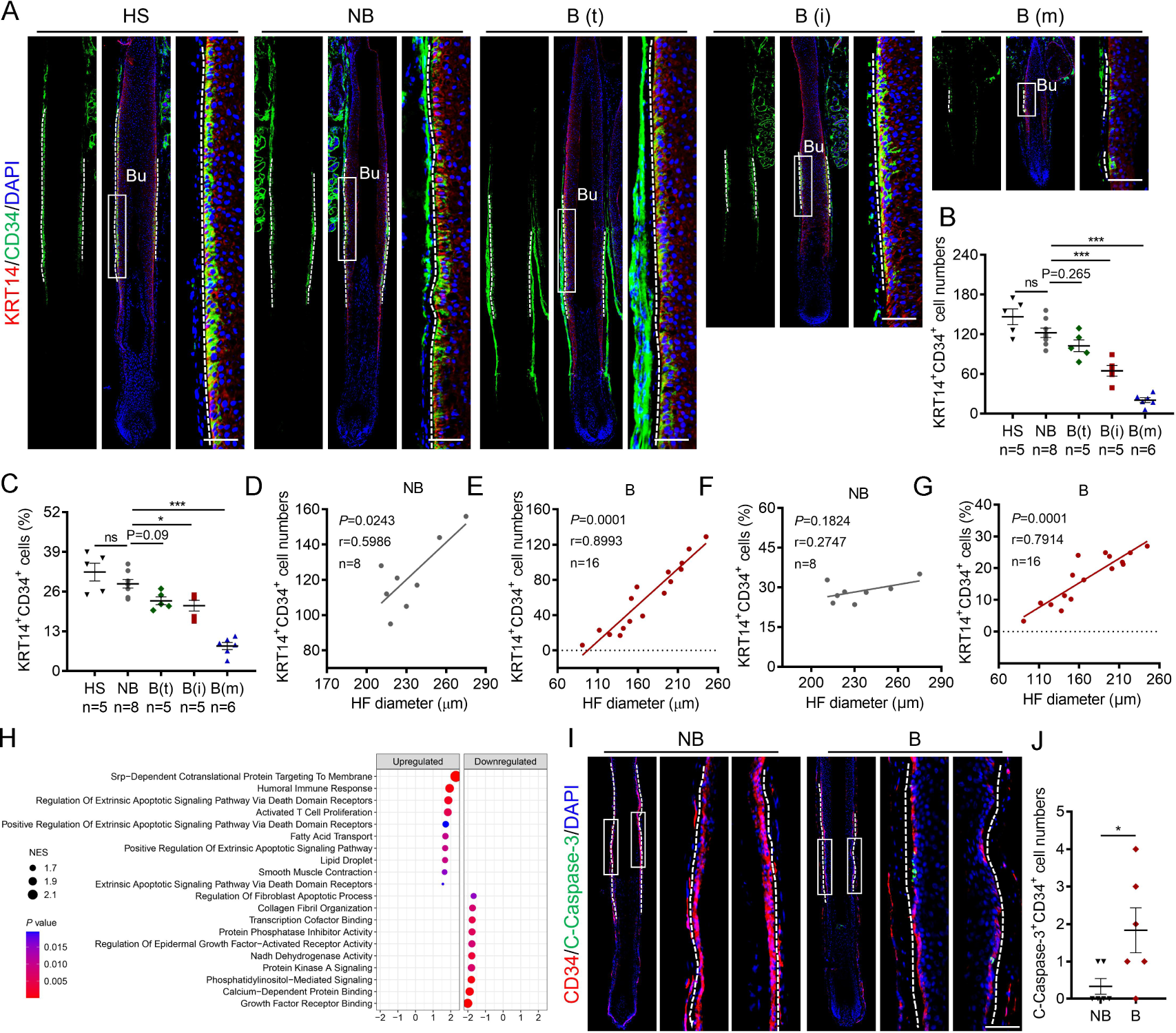
Apoptotic loss of HF progenitor cells is associated with HF miniaturization in AGA. **A** Immunostaining images show progenitor cells in HS terminal HFs (n=5 anagen HFs from 3 individuals), NB terminal HFs (n=8 anagen HFs) and B terminal /intermediate /mini HFs (B(t)/B(i)/B(m), n=5/5/6 anagen HFs) from 3 AGA patients. Epidermal cells and progenitor cells were labelled with KRT14 (red) and CD34 (green) respectively and ORS–DS boundary is indicated by dashed lines. Right panels, magnified images of boxed areas. Bu, bulge. Types of HFs were defined by their diameter. Terminal: diameter>200 μm; Intermediate: 150 μm <diameter<200 μm; mini: diameter<150 μm. **B, C** Quantification of the number of KRT14^+^CD34^+^ progenitor cells in different groups (**B**) and their proportion in the total outer layer of HFs (**C**). **D-G** Scatter plots show the correlation between HF diameters and the number or percentage of KRT14^+^CD34^+^ progenitor cells in AGA patients. Spearman’s correlation coefficient was used for the correlation analysis (two-tailed). **H** Top10 GO terms upregulated or downregulated in B versus in NB CD34^+^ progenitor cells. NES stands for enrichment scores. The color keys from red to blue indicate the range of P value. **I, J** Representative images of C-Caspase-3 and CD34 co-immunostaining on anagen HFs from NB and B scalps of AGA patients (n=6 HFs from 3 donors for each group); Right panels, magnified images of boxed areas (**I**). The quantification of C-Caspase-3^+^CD34^+^ cell numbers in two groups (**J**). Progenitor cells were indicated with dotted line. Scale bar: 50μm. The data represent the means±SEM. *P<0.05, **P<0.01, ***P<0.001, determined by one-way ANOVA with Tukey’s post hoc test (**B, C**) or unpaired Student’s t-test (**J**). ns, not significant.

Next, investigation into the causes of the progenitor cell loss via signaling pathway enrichment analysis revealed upregulated apoptotic signals in progenitor cells of balding anagen HFs (Fig. 3H). A significant increase in apoptosis was observed in the CD34^+^ progenitor cells of balding anagen HFs according to Cleaved Caspase-3 staining (Fig. 3I, J). These results support the notion that an apoptosis-induced gradual loss of CD34^+^ progenitor cells may be involved in the HF miniaturization in AGA.

### Hyperactive CTS contraction drives premature hair regression via inducing apoptosis of progenitor cells in AGA

Intimate crosstalk between stem cells and the surrounding microenvironment is vital for stem cell fate decisions. On the outside of the progenitor cells is a layer of elastic DS in murine pelage HF, while in human scalp HF is the multi-layer of connective tissue sheath (CTS) mainly composed of DS cells, collagens and vasculature (including vSMCs and endothelial cells) (*37*)(Fig. S6A; Fig. 4A, B). Pathway enrichment analysis revealed increased contractile activity in both DS cells and vSMCs of balding anagen HFs compared to non-balding or healthy groups (Fig. 2O and R), which can be confirmed by the immunostaining of phosphorylated myosin light chain 2 (p-MLC2), a marker of myosin active state(*40*) (Fig. 4C, D; Fig. S6B, C). Collectively, these results suggest that CTS contraction is enhanced in anagen HFs of balding scalps in AGA patients.

**Fig. 4:**
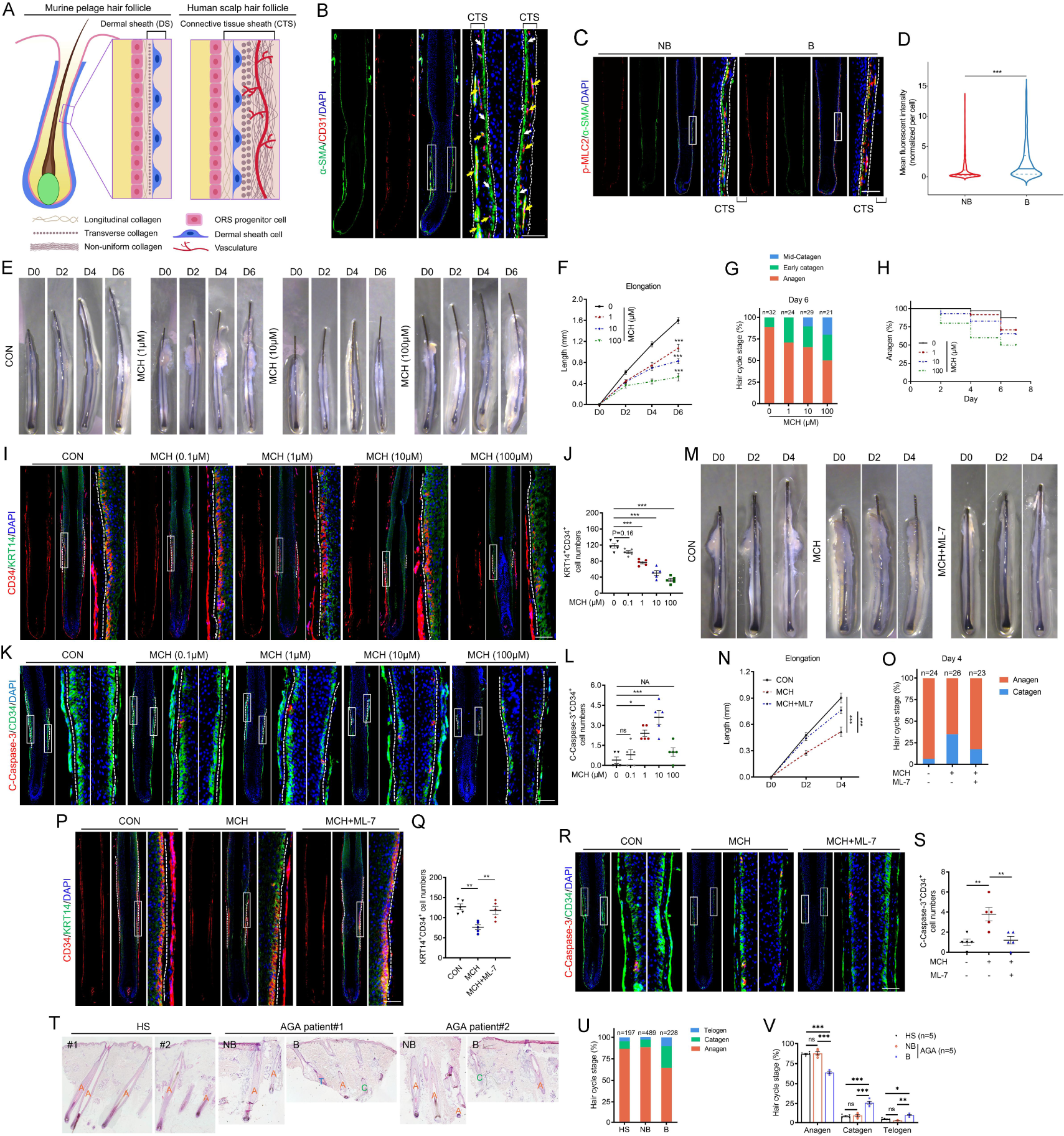
Hyperactive CTS contraction induces premature hair regression in AGA. **A** Dermal sheath (DS)/ connective tissue sheath (CTS) in murine pelage follicle/ human scalp follicle. This image is adapted from the previous publication (*37*). In murine pelage follicle (left), the DS is the outer layer of the ORS progenitors. In human scalp follicle, the outer layer is the CTS (right), which is mainly composed of DS cells, vasculature and multiple types of collagens. **B** Immunostaining of CD31 (an endothelial cell marker) and α-SMA (a marker for both DS and vSMC cells) in human scalps with anagen HFs. CTS is located between the two dotted lines. White arrows indicate DS. Yellow arrows indicate blood vessels. Right panels, magnified images of boxed areas. **C** The activation of CTS contraction was evaluated with p-MLC2 immunostaining on anagen HFs from non-balding occipital scalps (n=5 HFs) and balding frontal scalps (n=6 HFs) of 3 AGA patients. Right panels, magnified images of boxed areas. **D** Quantification of relative mean fluorescent intensity (RFI) of p-MLC2 in α-SMA positive cells in CTS in different groups. **E** Representative images of hair shaft elongation of human HFs ex vivo with vehicle (CON) or MCH treatment at concentration of 0.1 μM, 1 μM, 10 μM, 100 μM as indicated (n=21-32 HFs from 3 donors for each group). The same individuals’ HF samples were used in the following staining of HFs with the same treatments. **F** Quantification of hair shaft elongation on day 2/4/6 with indicated treatments (n=32/24/29/21 in different groups respectively). **G** Macroscopic quantification of hair cycle stage of HFs with vehicle or MCH treatments on day 6. **H** Percentages of HFs in anagen on day 2/4/6 with indicated treatments. **I** Representative images of co-immunostaining of CD34 and KRT14 in HFs with vehicle or MCH treatments on day 4. **J** Quantification of KRT14^+^CD34^+^ progenitor cell numbers in the total outer layer of HFs. **K** Apoptosis of progenitor cells were indicated with co-immunostaining of CD34 and C-Caspase-3 on day 2. Progenitor cells were indicated with and located inside the dotted lines. Right panels, magnified images of boxed areas. **L** Quantification of C-Caspase-3^+^CD34^+^ progenitor cells in HFs. To be mentioned, C-Caspase-3^+^CD34^+^ progenitor cells are barely detectable under 100 μM MCH administration because CD34+ progenitor cells were almost vanished under this condition. **M** Representative images of hair shaft elongation of human HFs ex vivo with vehicle or MCH (10 μM) and ML-7 (0.3 μM) treatment as indicated (n=23-26 HFs from 3 donors for each group, independent experiments). **N** Quantification of hair shaft elongation on day 2/4 with vehicle or MCH and ML-7 treatments (n=24/26/23 in different groups respectively). **O** Macroscopic quantification of hair cycle stage of HFs with vehicle or MCH and ML-7 treatments on day 4. **P** Co-immunostaining of CD34 and KRT14 on day 4. Right panels, magnified images of boxed areas. **Q** Quantification of total KRT14^+^CD34^+^ progenitor cell numbers in HFs (n=5 HFs from 3 donors for each group). **R** Co-immunostaining of CD34 and C-Caspase-3 showed progenitor cell apoptosis with indicated treatments on day 2 (n=5 HFs from 3 donors for each group). Right panels, magnified images of boxed areas. **S** Quantification of C-Caspase-3^+^CD34^+^ progenitor cells numbers in HFs (n=5 HFs from 3 donors for each group). **T** Representative images of HE staining with HS frontal scalps, NB occipital and B frontal scalps from AGA patients. A, anagen HF. C, catagen HF. T, telogen HF. **U** Macroscopic quantification of hair cycle stage of HFs in HE staining (n=197/489/228 HFs from 5 HS and 5 AGA donors). **V** Percentage of anagen, catagen and telogen HFs in HS, NB and B scalps (n=197/489/228 HFs from 5 HS and 5 AGA donors). Scale bar, 50 μm. Data are expressed as mean±SEM and were analyzed by one-way ANOVA with Tukey’s post hoc test (J, L, Q, S, V) and two-way ANOVA with a post hoc Holm– Sidak’s multiple comparisons test (F, N), or unpaired Student’s t-test (D). *p<0.05, **p<0.01, ***p<0.001. ns, not significant. NA, not applicable.

Given that the contractile force of DS is a key driver of HF regression during hair regeneration in mice(*12*), we wondered whether excessive contraction of the progenitor cell-encapsulating CTS, the analogous structure of murine DS in human, might affect hair follicle by modulating progenitor cell fate in AGA. To this end, we utilized methacholine (MCH)(*41, 42*) and norepinephrine (NE)(*43, 44*) to activate CTS contraction in the in vitro cultured human anagen HF organs. The efficiency of MCH and NE on contraction activation in this model was evaluated and verified by p-MLC2 immunostaining (Fig. S6D). Strikingly, HF growth was significantly inhibited (Fig. S6E-G) and premature catagen induction was promoted (Fig. S6H, I) in normal anagen HFs treated with MCH or NE. Furthermore, the observed activation of CTS contraction by MCH resulted in a dose-dependent negative effect on hair growth and anagen maintenance (Fig. 4E-H, and Fig. S7A, B). To figure out whether progenitor cells act as the targets of CTS contraction, we examined the effect of MCH on the number and apoptosis of HF progenitor cells. The immunostaining of CD34 and Cleaved Caspase-3 showed that activated CTS contraction by MCH promoted the loss and apoptosis of CD34^+^ progenitor cells in a concentration-dependent manner. Of note, Cleaved Caspase-3^+^CD34^+^ cells are barely detectable under 100 μM MCH treatment due to the fact that CD34^+^ progenitor cells were almost depleted under this condition (Fig. 4I-L).

Notably, blockade of CTS contraction with ML-7 (Fig. S7C), a specific myosin activation inhibitor(*45*), not only improved the growth retardation and catagen premature induction (Fig. 4M-O, and Fig. S7D), but also restored the apoptotic loss of progenitor cells in MCH-treated human HFs (Fig. 4P-S), confirming the critical role of CTS contraction in MCH effects. Consistently, we demonstrated that the percentage of catagen HFs is indeed markedly increased in the B scalps of AGA patients compared to NB or HS scalps (Fig. 4T-V). Collectively, these findings significantly strengthen the link between enhanced CTS contraction and AGA pathogenesis.

Subsequently, to determine whether apoptosis is necessary, we applied Z-VAD-FMK to block cell apoptosis(*46*). Our results showed that the observed phenotypes caused by MCH, including HF growth inhibition (Fig. 5A-D), premature catagen induction (Fig. 5E) and depletion of CD34^+^ progenitor cells (Fig. 5F-I), were counteracted after Z-VAD-FMK administration. Taken together, these data suggest that over-activated CTS contraction compresses hair growth and drives premature hair regression via promoting the apoptosis of progenitor cells.

**Fig. 5:**
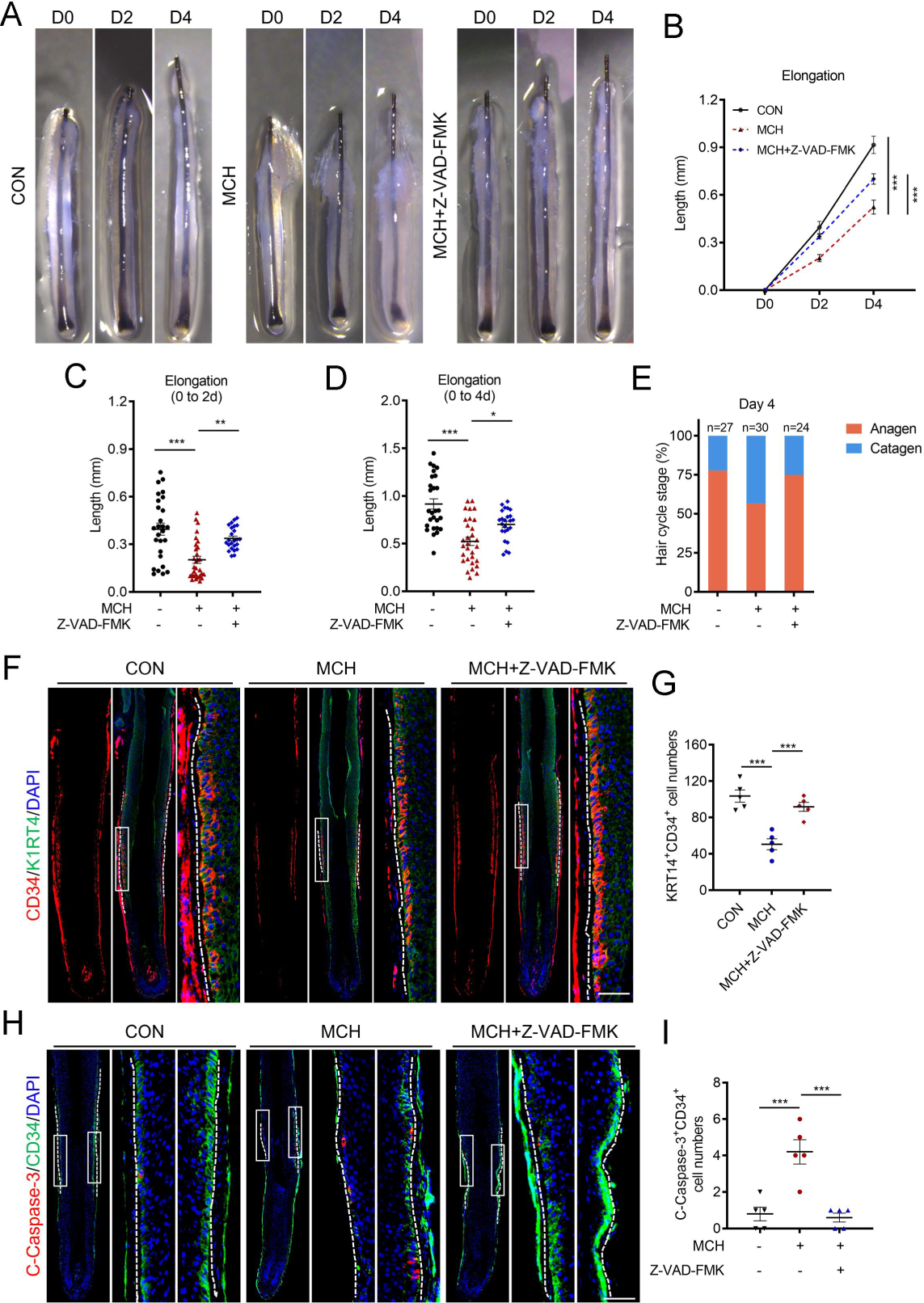
CTS contraction drives premature hair regression in an apoptosis-dependent manner. **A** Representative images of hair shaft elongation of human HFs ex vivo with or without MCH (10 μM) and Z-VAD-FMK (10 μM) as indicated (n=24-30 anagen HFs from 3 donors for each group). The same individuals’ HF samples were used in the following staining of HFs with the same treatments. **B** Quantification of hair shaft elongation on day 2/4/6 with indicated treatments. **C, D** Scatter plots show the elongation length of hair shaft with indicated treatments on day 2 (**C**) and day 4 (**D**). **E** Macroscopic quantification of hair cycle stage of HFs with indicated treatments on day 4. **F** Co-immunostaining of CD34 and KRT14 on day 4. Right panels, magnified images of boxed areas. **G** Quantification of total KRT14^+^CD34^+^ progenitor cell numbers in HFs with indicated treatments (n=5 HFs for each group). **H** Co-immunostaining of CD34 and C-Caspase-3 showed apoptosis of progenitor cells with indicated treatments on day 2 (n=5 HFs for each group). **I** Quantification of C-Caspase-3^+^CD34^+^ progenitor cell numbers in HFs with indicated treatments. Progenitor cells were indicated with dotted line. Right panels, magnified images of boxed areas. Scale bar: 50 μm. The data represent the means±SEM. *p<0.05, **p<0.01, ***p<0.001, determined by one-way ANOVA with Tukey’s post hoc test (C, D, G, I) and two-way ANOVA with a post hoc Holm–Sidak’s multiple comparisons test (B).

### CTS contraction induces progenitor cell apoptosis and premature hair regression via PIEZO1 signaling in AGA

To pinpoint the specific molecular mechanisms by which the contractility of CTS regulates progenitor cells, we first examined the expression levels of the known mechanosensitive ion channel protein PIEZO1, which has been found to be highly expressed in HFSCs in mice(*47*). Our results showed that PIEZO1 was abundantly expressed in CD34^+^ progenitor cells (Fig. S8A) and was expressed at much higher levels in balding anagen HFs compared to non-balding group (Fig. 6A, B). We then used intracellular Ca^2+^ as a downstream indicator of PIEZO1 signaling(*48, 49*). As quantified by Fluo-4 (a Calcium influx indicator) RFI, there was a significant increase in intracellular Ca^2+^ concentration in CD34^+^ progenitor cells of balding anagen HFs (Fig. 6C, D). These results suggest that PIEZO1 signaling is activated in progenitor cells of balding anagen HFs.

**Fig. 6:**
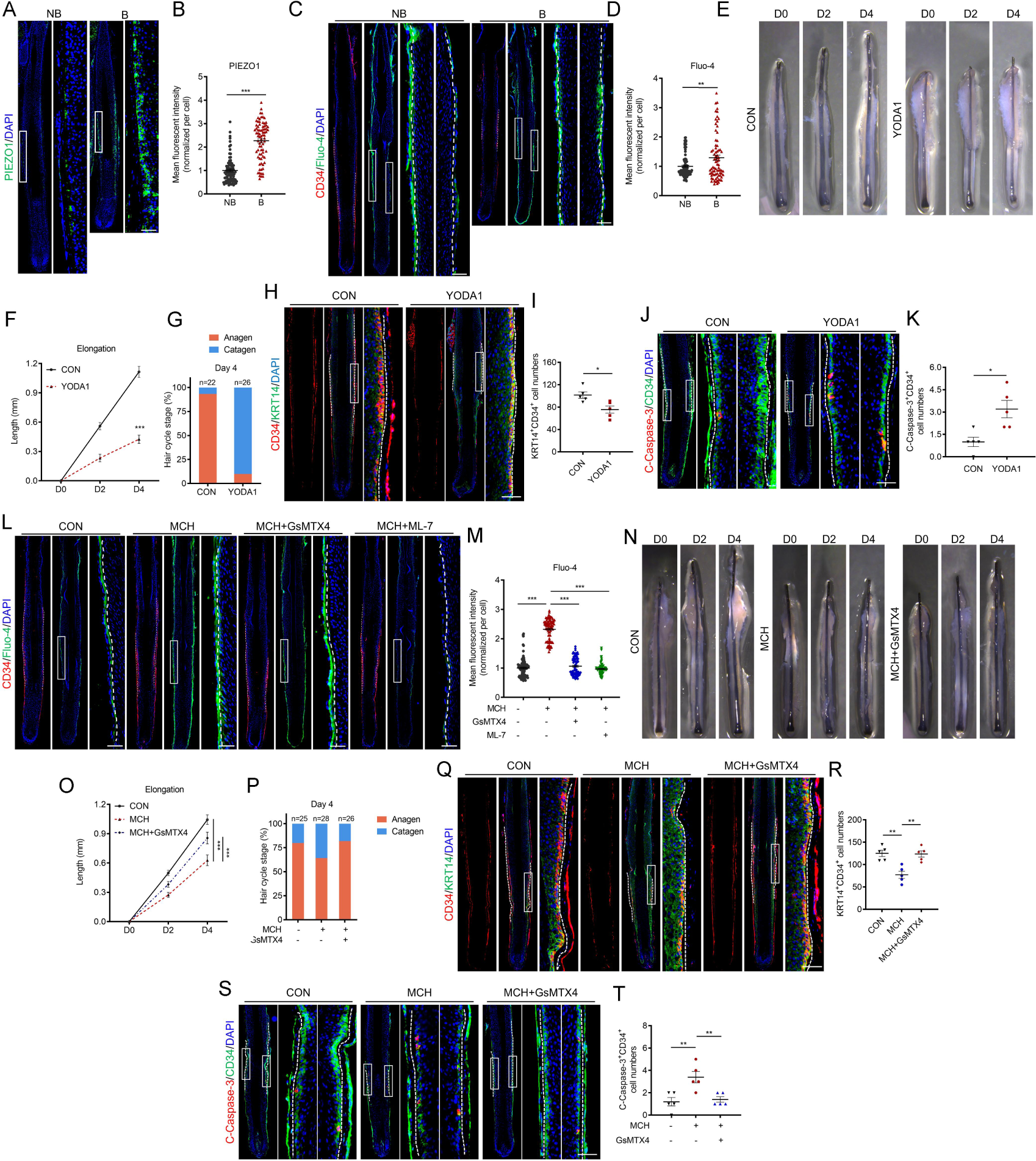
PIEZO1 signaling is necessary for CTS contraction-induced progenitor cell apoptosis and premature hair regression. **A, B** (**A**) Representative immunostaining of PIEZO1 on skin sections from balding frontal and non-balding occipital scalps of AGA patients (n=3) and (**B**) the quantification of mean fluorescent intensity in outermost layer of mid-HF cells (n=70-90 cells from 3 patients). **C, D** (**C**) CD34 and Fluo-4 staining were performed on serial sections from balding frontal and non-balding occipital scalps of AGA patients (n=3) and (**D**) the Fluo-4 mean fluorescent intensity in CD34^+^ progenitor cells was quantified (n=100 cells in non-balding HFs and 80 cells in balding HFs from 3 patients). Fluo-4 indicates the cellular Ca^2+^ concentration. **E** Representative images of hair shaft elongation of human HFs ex vivo with or without YODA1 (15 μM) as indicated (n=22-26 anagen HFs from 3 individuals for each group). The same individuals’ HF samples were used in the following staining of HFs with the same treatments. **F** Quantification of hair shaft elongation on day 2/4 with vehicle or YODA1 treatments. **G** Macroscopic quantification of hair cycle stage of HFs with YODA1 treatment on day 4. **H** Co-immunostaining of CD34 and KRT14 on day 4. **I** Quantification of total KRT14^+^CD34^+^ progenitor cell numbers (n=5). **J** Co-immunostaining of CD34 and C-Caspase-3 showed apoptosis of progenitor cells on day 2. **K** Quantification of C-Caspase-3^+^CD34^+^ apoptotic progenitor cell numbers in HFs (n=5). **L, M** (**L**) CD34 and Fluo-4 staining were performed on serial sections of HFs with indicated treatments (n=3 HFs for each group) and (**M**) the Fluo-4 mean fluorescent intensity in CD34^+^ progenitor cells was quantified (n=80-100 cells from 3 different HFs). **N** Representative images of hair shaft elongation of human anagen HFs ex vivo treated with MCH (10 μM) and GsMTX4(1 μM) (n=25 HFs for CON, 28 HFs for MCH and 26 HFs for MCH+GsMTX4; from 3 individuals). The same individuals’ HF samples were used in the following staining of HFs with the same treatments. **O** Quantification of hair shaft elongation on day 2/4 with MCH and GsMTX4 treatments. **P** Macroscopic quantification of hair cycle stage of HFs with MCH and GsMTX4 treatments on day 4. **Q** Co-immunostaining of CD34 and KRT14 on day 4. **R** Quantification of total KRT14^+^CD34^+^ progenitor cell numbers in HFs (n=5 for each group). **S** Co-immunostaining of CD34 and C-Caspase-3 showed apoptosis of progenitor cells on day 2. **T** Quantification of C-Caspase-3^+^CD34^+^ progenitor cell numbers in HFs (n=5 for each group). Right panels, magnified images of boxed areas. Scale bar: 50 μm. Data are expressed as mean±SEM and were analyzed by one-way ANOVA with Tukey’s post hoc test (M, R, T), two-way ANOVA with a post hoc Holm–Sidak’s multiple comparisons test (F, O), and two-tailed unpaired Student’s t-test (B, D, I, K). *p<0.05, **p<0.01, ***p<0.001.

We next investigated whether activation of PIEZO1 is sufficient to induce HF growth retardation and premature regression. To this end, we applied YODA1, a PIEZO1-specific activator to cultured anagen HFs (Fig. S8B, C). Intriguingly, anagen HFs treated with YODA1 displayed a phenotype similar to MCH (Fig. 6E-K and Fig. S8D). In addition, the Ca^2+^ concentration in MCH-treated HF progenitor cells was elevated and could be reversed by the specific PIEZO1 inhibitor GsMTX4 and the myosin activation inhibitor ML-7 (Fig. 6L, M), indicating the activation of PIEZO1 signaling by CTS contraction. Not surprisingly, inhibition of PIEZO1 signaling by GsMTX4 rescued the phenotypes induced by MCH (Fig. 6N-T and Fig. S8E).

These results prompt us to speculate that CTS contraction induced in balding anagen HFs activates the PIEZO1 signaling pathway in progenitor cells via mechano-transduction, thereby inducing progenitor cell apoptosis and thus promoting premature HF regression.

### Targeting CTS contraction by ML-7 improves the growth of HFs from the balding scalps of AGA patients

All the above results inspired us to wonder whether targeting CTS contraction is a promising therapy for AGA. We first obtained anagen HFs from the balding scalps of AGA patients, which had been experiencing the process of miniaturization. As expected, the balding anagen HFs exhibited slower growth compared with those from occipital non-balding scalps of the same AGA patients in ex vivo organ culture system (Fig. S9A-C). We then treated the balding HFs with ML-7, or minoxidil (MNX), the FDA-approved drug for AGA treatment. Immunostaining of p-MLC2 confirmed that ML-7 abolished muscle contraction activity in CTS, but MNX did not (Fig. S9D). Excitingly, ML-7 could significantly improve the growth and premature regression of balding HFs of AGA patients, even better than MNX (Fig. 7A-D; Fig. S9E). Consistently, ML-7 showed a better effect on decreasing the apoptotic loss of progenitor cells compared with MNX (Fig. 7E, F). Furthermore, we demonstrated ML-7 did not affect the growth and anagen-to-catagen progression of cultured anagen HFs from occipital non-balding scalps of AGA patients, suggesting a specific therapeutic effect for balding anagen HFs with enhanced CTS contraction (Fig. S9F-H). Taken together, these findings suggest that relaxing HFs via targeting CTS contraction is a hopeful strategy for AGA therapy.

**Fig. 7:**
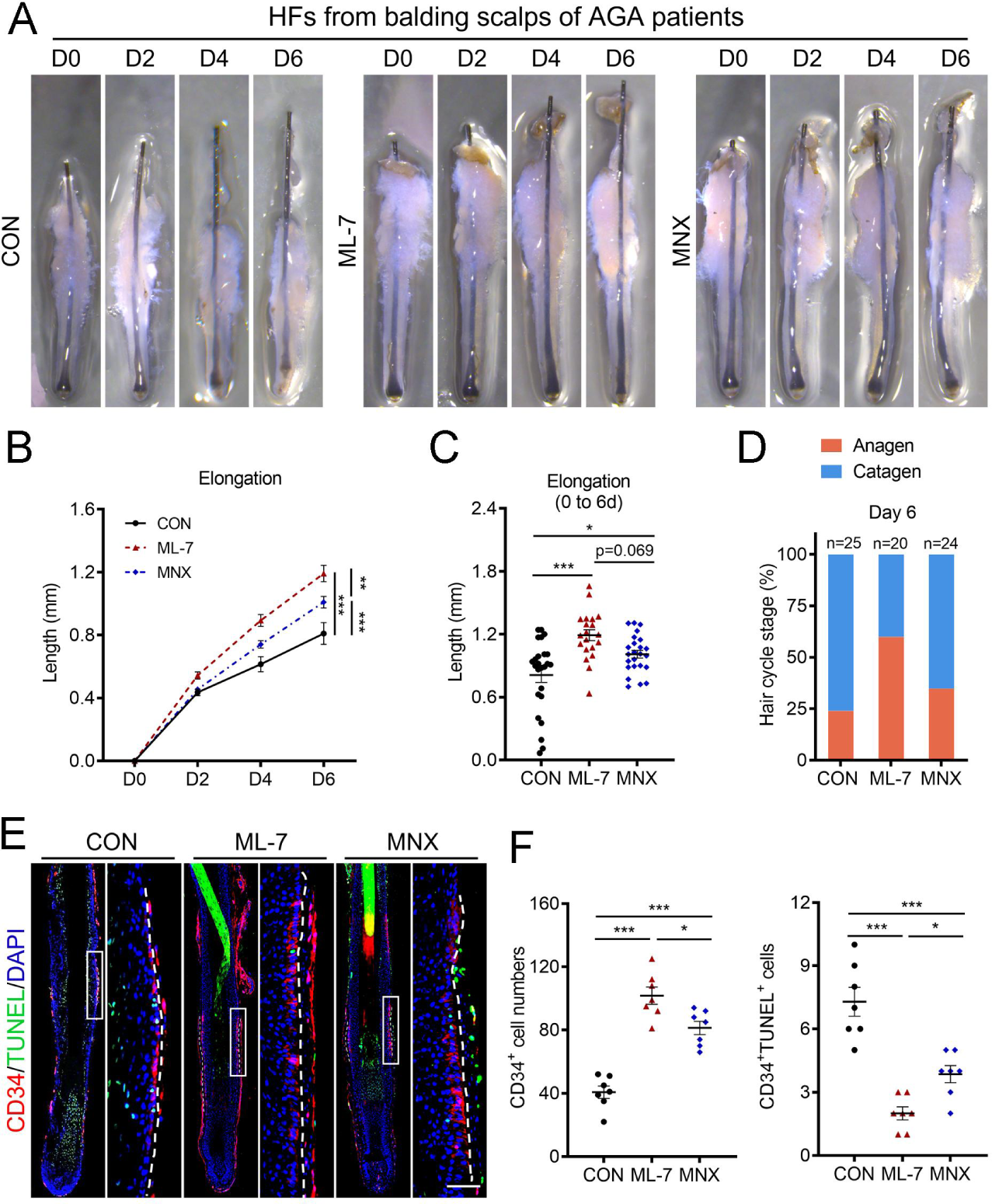
ML-7 exhibits a better effect than MNX on promoting the growth of HFs from the balding scalps of AGA patients. **A** Representative images of hair shaft elongation of anagen HFs from balding scalps of AGA patients ex vivo treated with vehicle (CON), ML-7 (0.3 μM) or minoxidil (MNX) ((100 μM)) as indicated (n=31-37 balding HFs from 5 AGA patients for each group). 3 patients’ HF samples were used in the following staining. **B** Quantification of hair shaft elongation on day 2/4/6 with indicated treatments. **C** Scatter plots show the elongation length of hair shaft on day 6. **D** Macroscopic quantification of hair cycle stage of HFs with indicated treatments on day 6. **E** Co-immunostaining of CD34 and TUNEL showed apoptosis of progenitor cells with indicated treatments (n=7 balding HFs from 3 AGA patients for each group). Right panels, magnified images of boxed areas. **F** Quantification of total and apoptosis CD34^+^ progenitor cell numbers in HFs with indicated treatments. Scale bar: 50 μm. The data represent the means±SEM. *p<0.05, **p<0.01, ***p<0.001, determined by one-way ANOVA with Tukey’s post hoc test (C, F) and two-way ANOVA with a post hoc Holm–Sidak’s multiple comparisons test (B).

## Discussion

Here we have presented what is, to the best of our knowledge, the first and currently the only comprehensive single-cell transcriptome profiling of all cell types within HF units from balding and non-balding scalps of AGA patients compared with healthy individuals. Considering that HFs of balding scalps used in the present study were in anagen phase and at the early stage of hair miniaturization, isolated from the edge of frontal hairline of AGA patients, our high-quality data allowed us to illustrate the early changes in cell subpopulations, the HF cell lineage trajectory and cell-cell communications in AGA in unprecedented detail. Our atlas uncovers at least 16 different broad cell types in human HF unit, which are more than those described in previous scRNA-seq studies(*25, 26*). The reason for this obvious difference could be likely attributed to the optimization of human HF single cell isolation by combining mechanical separation with multienzyme digestion. Based on this, we obtained previously undescribed DP cells and major cell types of CTS (including DS, vSMC and endothelial cells) in human HFs, which are reported to be essential for regulating the growth of hair follicles(*12, 37, 50*). Delineation of DP and CTS cells at single-cell level provides an opportunity to determine the precise roles of these cells in the pathogenesis of AGA.

The hair growth is powered by HFSCs (termed as bulge stem cells) that have the ability to give rise to all epithelial components of the hair follicle(*51*). However, current understanding of the cellular and molecular changes in the bulge in AGA development is limited. Our scRNA-seq data show that bulge stem cells contain two major subclusters: CD200^+^ bulge stem cells and CD34^+^ progenitor cells. Thereinto, the number and percentage of progenitor cells are significantly decreased in the balding anagen HFs of AGA patients. These results are consistent with the previous study showing that balding scalp in men with AGA retains HFSCs but lacks HF progenitor cells(*14*). Besides, we demonstrate that HFSCs are more likely to differentiate into interfollicular epidermal and sebaceous lineages, rather than HF lineage cells, in the balding scalps, which may be responsible for the progressive hair follicle miniaturization and hair thinning in AGA. Similarly, in the case of ageing- and obesity-induced hair loss in mice, those aberrant fate changes also occur in a fraction of HFSCs, which prefer to undergo epidermal and sebocyte differentiation regulated by COL17A1 proteolysis and inflammatory signals respectively, thereby decreasing the HFSC pool and leading to hair follicle miniaturization in a stepwise manner(*38, 39*). Here, our data also suggest an important role of COL17A1 in the aberrant differentiation of HFSCs in AGA development. During normal hair growth, HF stem/progenitor cells proliferate and differentiate to generate hair matrix cells, which rapidly proliferate to produce the hair shaft of growing hair in the end(*52*). Here, we show that the proliferating matrix cells are also decreased in balding HFs in AGA. It is tempting to speculate that the apoptotic loss of progenitor cells contributes to a decrease in matrix cells, which may cause the premature induction of catagen (namely shortening of anagen), and eventually weakens the generation of terminal hair shaft in AGA development.

DP acts as the signaling center to govern the proliferation and differentiation of HF stem/progenitor cells in hair growth(*11, 53*). Our previous study demonstrated that androgen receptor (AR) is upregulated in DP cells, which induce apoptosis of microvascular endothelial cells via paracrine signaling in the DP of balding HFs in AGA(*6*). However, in this study, except for the downregulated angiogenesis, our scRNA-seq analysis identifies no other obvious changes in cell subpopulations and signaling pathways, which may be involved in the regulation of hair growth, in DP cells of balding HFs from AGA patients. The reasons for this discrepancy are currently not clear, and further analysis and experimentation will be required to address this point. Although microinflammation has been commonly reported in AGA(*54–57*), the details remain largely undefined. Here, we identity that T cells are the main lymphocytes in human HF units, and AGA is featured by increased Th17 cells. Although a previous study has identified that progenitor cells are significantly diminished by flow cytometry in the bald scalp of AGA patients(*14*), the details in this aspect are obscure. In this study, we further confirm that CD34^+^ progenitor cells are decreased in the balding anagen HFs in AGA, and determine that these progenitor cells are derived from CD200^+^ bulge stem cells via scRNA-seq analysis and immunostaining. Importantly, we also find that the reduction in HF progenitor cells is attributed to the increased apoptosis, and the number and percentage of these cells is positively correlated with the hair follicle size in the balding scalp in AGA. These findings suggest a pivotal role for progenitor cell depletion in the pathogenesis of AGA.

HF growth is tightly spatiotemporally regulated by the crosstalk between stem/progenitor cells and the microenvironment, mainly the mesenchymal niche(*10*). In addition to DP whose role is well recognized, DS is also an important component of the mesenchymal niche. DS is continuous with the DP and generates an envelope surrounding the hair follicle, whose role has been far less examined than the DP yet may be equivalently important for the regulation of hair growth(*37*). Until recently, DS has been identified as smooth muscle that drives hair follicle regression for reuniting HFSCs and niche to regenerate tissue structure during hair regeneration in mice(*12*). Further study demonstrates that progenitor cell-derived endothelin governs DS contraction in mice(*58*). However, the role of CTS (the analogous structure of murine DS in human), in human hair follicle, especially in hair disorders, remains undefined. In this study, via single-cell sequencing analysis we identify that smooth muscle contraction activity is markedly increased in the major cell types of CTS, including DS and vSMC cells in balding anagen HFs of AGA patients. We further, by co-immunostaining, confirm this conclusion, and demonstrate that muscle contraction-related pathways are hyperactivated in the CTS of balding anagen HFs; In human hair follicle organ culture system, activation of muscle contraction in CTS drives premature hair follicle regression via inducing progenitor cell apoptosis. Surprisingly, we find that pharmacological inhibition of muscle contraction can obviously improve the growth and premature catagen induction of hair follicles from the balding scalps of AGA patients, even better than the FDA-approved drug minoxidil, strongly suggesting that targeting muscle contraction in CTS is a very promising therapy for AGA. It will also be interesting in future studies to determine the mechanism that hyperactivates the muscle contraction of CTS cells in the development of AGA.

Our results suggest PIEZO1, a mechanosensitive ion channel, as a regulator in HF progenitor cells that senses the mechanical force generated by CTS contraction. We show that PIEZO1 is expressed in HF progenitor cells, and it becomes markedly upregulated and activated in the balding anagen HFs of AGA patients. Inhibition of PIEZO1 signaling evidently alleviates progenitor cell apoptosis and premature hair follicle regression induced by CTS contraction; in contrast, PIEZO1 hyperactivation induces apoptosis of progenitor cells and premature entry into regression in human anagen hair follicles. Similarly, PIEZO1 has been reported to be abundantly expressed in HFSCs, and is responsible for niche-size-decrease-triggered ectopic HFSC apoptosis in mice(*47*). We conclude that the role of PIEZO1 in sensing the abnormal mechanical force from the microenvironment to regulate the HFSC apoptosis is conserved across species, and the mechanosensitive channel might be a therapeutic target for hair loss. However, the detailed molecular mechanisms by which aberrant PIEZO1 activation induces progenitor cell apoptosis in AGA need further study to clarify.

In summary, the present study reveals the cellular hierarchies via scRNA-seq analysis and demonstrates hyperactive CTS contraction as a critical cause of hair regression in AGA, strongly suggesting that relaxing hair follicle by targeting CTS might be a promising therapy for this disorder.

## Methods

### Human scalp and HF samples

Human scalp skin biopsies and HF units were collected from non-balding occipital and balding frontal sites of male AGA patients undergoing hair transplantation surgery, and normal frontal sites of male healthy volunteers from the Department of Dermatology in Xiangya Hospital, Central South University. The procedures were in accordance with protocols set out in the WMA Declaration of Helsinki and the Department of Health and Human Services Belmont Report and approved by the ethical committee of the Xiangya Hospital of Central South University and written informed consent was acquired from all participants.

### Identification of hair cycle phases in human HFs

Hair cycle phases of human HFs were determined by morphological analysis, HE staining, co-immunostaining of Ki67 and TUNEL, and immunostaining of Cleaved Caspase-3 (C-Caspase-3) as previously described(*22, 23*). With these identification methods (Fig. S1), all human HFs used in this study (including scRNA-seq, staining and in vitro organ culture) had been confirmed in the typical anagen phase before experimentation unless otherwise stated.

### Preparation of single-cell suspensions of HF units

To obtain single-cell suspension of entire HF units, DPs were isolated first by micro-dissection from HFs and were chopped. The remaining hair follicle tissue samples were cut with scissors and digested with the chopped DPs in 4mg/ml Dispase II and Collagenase IV for 40 min at 37°C, centrifuged at 200g for 5min. The pellet was resuspended and digested in 0.25% trypsin for 10 min at 37°C, washed in PBS+5%FBS and filtered through a 70 μm cell strainer. The Dead Cell Removal Kit (Miltenyi Biotec) was used in accordance with the manufacturer’s instruction to obtain live cells. 10 μL of the cell suspension were mixed with the same volume of Trypan blue for cell counting. The loading volume for sequencing was verified by the cell concentration

### ScRNA-seq

scRNA-seq mainly includes GEM (gel bead-in-emulsion) generation, barcoding, cDNA amplification, library construction and sequencing. These steps were completed according to the user’s instructions of Chromium Single Cell 3ʹ Reagent Kits v3.1 (10× Genomics, product code: 1000268, 1000215, 1000120) (https://www.10xgenomics.com/support/single-cell-gene-expression). Libraries were sequenced by an Illumina NovaSeq6000 System. Approximately 10,000 cells (targeting 5000–12,000) per sample were used for single-cell RNA sequencing.

### ScRNA-seq data processing and cell type identification

We used the default parameters of Cell Ranger software suite (v6.0.2) to align and quantify the raw reads data from 10x Genomics. After the initial Cell Ranger metric assessment, Seurat (v4.3.0)(*59*) in R version 4.3.1 was used to exclude cells with fewer than 200 genes or more than 6000 genes detected, and more than 20% mitochondrial reads for downstream analysis.

After quality control, 76368 cells remained and were used for downstream bioinformatic analysis. The “NormalizeData” function was used to normalize the feature expression measurements for each cell by the total expression. The “ScaleData” function was used to scale and center the expression of each gene for dimensional reduction. To avoid batch effects among samples and experiments, we integrated data from all samples using harmony (v0.1.1). Total cell clustering was performed by “FindClusters”function with Louvain algorithm at a resolution of 0.2. Non-linear dimensional reduction was performed by “RunUMAP” function and visualized by Uniform Manifold Approximation and Projection (UMAP). Markers genes of each cell cluster were determined by “FindAllMarkers” function with Wilcoxon rank-sum test. Only those with |‘avg_logFC’| ≥0.25 and ‘adjusted P value’≤0.05 were considered as marker genes.

For subgroup cell clustering, cells of each cell type were extracted separately and clustered by their first 20 PCs and appropriate resolution. To further approximate the low-dimensional data manifold representing the differentiation trajectory for Bulge cells, spliced status of mRNAs in single-cell RNA sequencing data were used to estimations of RNA velocities through velocyto(*60*). Markers genes of each subcluster were identified by “FindAllMarkers” function with the default parameters. “DoHeatmap” function were used to show top marker genes in heatmap.

### Identification of balding-associated DEGs

We used the function of “FindMakers” in Seurat to identify balding-vs-nonbalding and balding-vs-healthy differentially expressed genes (DEGs) for each cell type and sub-cell type. The adjusted P values of each DEG were calculated by non-parametric two-sided Wilcoxon rank-sum test and only those with |‘avg_logFC’| > 0.25 and ‘adjusted P value’ < 0.05 were considered to be balding-associated DEGs.

### Function enrichment analysis

Gene Set Enrichment Analysis (GSEA) of GO and KEGG was performed by clusterProfiler R package (v4.5.1.902) and visualized with ggplot2 R package (v3.4.1)(*61*). Representative terms selected from the top ranked GO terms and KEGG pathways from MsigDB were displayed.

### Pseudotime analysis

Monocle3 was used to reconstruct the differentiation trajectory for all cell types(*36*). The expression matrix of all cells was used as input and the UMAP embedding coordinates were import from Seurat result. The start points of bulge and IFE basal cells were specified as the root nodes separately for the SOX9^+^ HF branch and SOX9^-^ IFE branch of the trajectory graph by “order_cells” function. “Plot_cells” function was used to visualize each cell type along the pseudotime trajectory. To further reconstruct the differentiation trajectory and do “branched expression analysis modeling” (BAEM) for bulge cell fate cell determination related cell types, such as bulge to ORS and IFE, Monocle2(*62*) was used to reconstruct the differentiation trajectory for small set of cell types and “BAEM” function were performed to identify cell determination related molecules.

### Cell-cell communication analysis

To assess cell-cell communications among different cell types, we used cellchat (v1.1.3) to infer the intercellular communication network from single-cell RNA-seq data. Only cell types with more than 10 cells were considered in the analysis. The “trimean” method is used for calculating the average gene expression per cell group in “computeCommunProb” function. Pairwise comparison and visualization are performed by cellchat build-in function, only interactions with P value lower than 0.05 are considered to be real.

### HF organ culture

Human HFs were obtained from the non-balding occipital sites (unless stated otherwise) of male AGA patients undergoing hair transplantation surgery. After microdissection, the HFs were first incubated in William’E medium (WEM, ThermoFisher) for 24 hours to re-equilibrate. HFs in anagen VI phase were randomly assigned to the different experimental groups. HFs were cultured at 37°C with 5% CO_2_ in WEM supplemented with 2 mM of L-glutamine, 10 ng/ml hydrocortisone, 10 ug/ml insulin and 1% penicillin/streptomycin mixure, and photographed under a stereoscope every two days to count hair shaft length and hair cycle stage. The hair cycle stage (anagen/early-catagen/mid-catagen) of HFs was determined according to the identification methods described above(*22, 23*).

### Chemical stimulation of HFs

All the chemicals were purchased from Selleck. After 24 hours, medium was replaced and HFs were treated with vehicle (0.1% DMSO), MCH (0.1,1 10,100 μM as indicated), NE (0.1 μM), ML-7(0.3 μM), Z-VAD-FMK (10 μM), YODA1(15 μM), or GsMTX4(1 μM) according to the design of different experiments and the medium was changed daily with the indicated treatment. HFs were cultured for 2-10 days for analysis of hair growth and immunostaining.

### Immunostaining and TUNEL staining

Immunostaining was performed as previously described(*63*). Briefly, OCT-embedded samples were sectioned (10 μm) with a Leica cryostat. The sections were fixed for 10 min with 4% PFA, and washed with PBS, then blocked for 60 min with blocking buffer (5% NDS, 1% BSA, 0.3% Triton X-100). Primary antibodies were incubated overnight at 4°C. Secondary antibody was incubated for 60 min at room temperature. After washed, sections were counterstained with DAPI. TUENL staining was performed with TUNEL assay kit (Roche, USA) according to manufacturer’s instructions, and then stained with CD34 following the immunostaining protocol. For the co-immunostaining of two antibodies from the same host species, double immunohistochemistry for paraffin sections (8 μm) was conducted by using the Opal 4 color manual immunohistochemistry (IHC) kit (NEL810001KT, PerkinElmer). The signal of p-MLC2 was amplified by the Opal 4 color manual IHC kit (NEL810001KT, PerkinElmer). All pictures were taken with a Zeiss Axioplan 2 microscope. The fluorescence intensity was evaluated with ImageJ. The primary antibodies were used in this study: Rabbit anti-SOX9 (1:100, Cell Signaling), Mouse anti-KRT15 (1:2000, Lab vision), Mouse anti-KRT14 (1:4000, Abcam), Rabbit anti-KRT6A (1:500, Atlas Antibodies), Rabbit anti-KRT10 (1:1000, Abcam), Mouse anti-α-SMA (1:4000, Abcam), Rabbit anti-CD200 (1:200, Cell Signaling), Rabbit anti-CD34 (1:250, Abcam), Rabbit anti-Cleaved Caspase-3 (1:400, Cell Signaling), Rabbit anti-p-MLC2 (1:100, Cell Signaling), Rabbit anti-Ki67 (1:500, Abcam), Rabbit anti-PIEZO1 (1:100, Proteintech). Staining intensity was evaluated in well-defined reference by quantitative (immuno-) histomorphometry using NIH ImageJ software.

### In vitro Ca^2+^ imaging

HFs were incubated with Fluo-4 staining solution (Beyotime) in culture medium at 37°C for 1 hour. Afterward HFs were washed with DPBS and cultured DPBS with 1mM calcium. For different treatments, MCH, ML-7, YODA1 or GsMTX4 was added to the culture medium for 30 min. The HFs were embedded in OCT and were sectioned with a Leica cryostat. Images were taken immediately to record the fluorescence intensity to indicate Ca^2+^ concentration and the adjacent section was used for CD34 staining to indicate progenitor cells.

### Statistical analysis

All data were expressed as mean±SEM and were analyzed by ANOVA or Kruskall-Wallis test when more than two groups were compared or Student’s t-test when only two groups were compared. Graphpad Prism 8, R (v4.1.3) and Excel (Microsoft) were used to assess statistical significance. Statistical significance was set at a P < 0.05. *, ** and *** indicate P < 0.05, P < 0.01 and P < 0.001, respectively.

## Data availability

All data needed to assess the conclusions in the present study are provided in the manuscript and/or the Supplementary Materials. The scRNA-seq datasets are available from the genome sequence archive under accession number HRA005629 (http://bigd.big.ac.cn/gsa-human/). Any other data supporting the findings of this study are available from the corresponding author upon reasonable request.

## Acknowledgements

This work was supported by the National Natural Science Funds for Distinguished Young Scholars (No. 82225039), the National Key Research and Development Program of China (No. 2023YFC2509003), the National Natural Science Foundation of China (No. 82304057, No. 81874251, No. 82073457, No. 82173448, No. 82373508, No. 82303992), the Natural Science Funds of Hunan province for excellent Young Scholars (No. 2023JJ20094), the Natural Science Foundation of Hunan Province, China (No. 2021JJ31079). We thank our colleagues (Department of Dermatology, Xiangya Hospital, Central South University, China) for their generous support throughout this work.

## Author contributions

J.L., Z.D., G.L. and L.Y. designed and conceived the study. Z.D., G.L. and L.Y. performed data analyses. L.Y. and Z.D. performed scRNA-seq analysis. G.L., Z.D., S.D. and M.C. performed most experiments. Y.Z., F.L., Y.T., Y.W., S.X., Z.W., B.W., Z.Z. and W.S. contributed to sample collection. Z.D. and M.C. help to generate sequencing libraries. H.X. and Y.Z. provide critical discussion and suggestion. Z.D., G.L., L.Y. and J.L. prepared the manuscript with input from coauthors.

## Competing interests

The authors declare no competing interests.

## Notes

### Competing Interest Statement

The authors have declared no competing interest.

### Summary of Updates

Figure 3, Figure 4, Figure 5, Figure 6 are revised, and Supplementary figures are updated.

