## Supplementary Figure 1-9 for "Single-cell transcriptomics reveals cellular hierarchies and aberrant CTS contraction-mediated premature hair regression in androgenetic alopecia"

### Supplementary Figures and figure legends

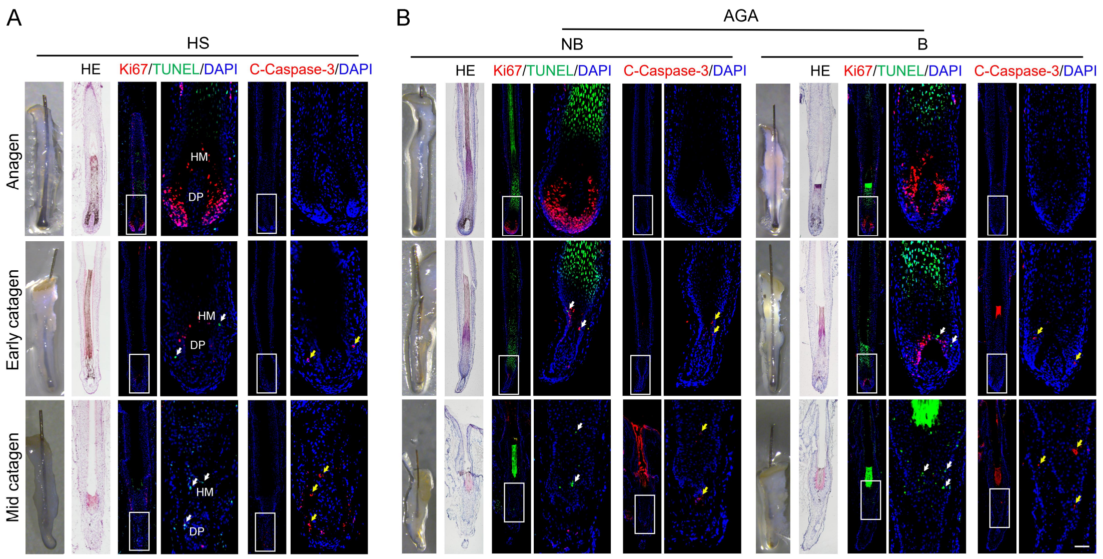


**Fig. S1: Identification of hair cycle phases of human HFs.**

**A, B** Representative images of morphological analysis, HE staining, co-immunostaining of Ki67 and TUNEL, and immunostaining of C-Caspase-3 for anagen, early catagen and mid catagen HS HFs (**A**), NB and B HFs from AGA patients (**B**). White arrows indicate TUENL positive cells. Yellow arrows indicate C-Caspase-3 positive cells. Right panels, magnified images of boxed areas. HM, hair matrix. DP, dermal papilla. Catagen HFs displayed obvious apoptotic signals in HM. Scale bar: 50 µm.

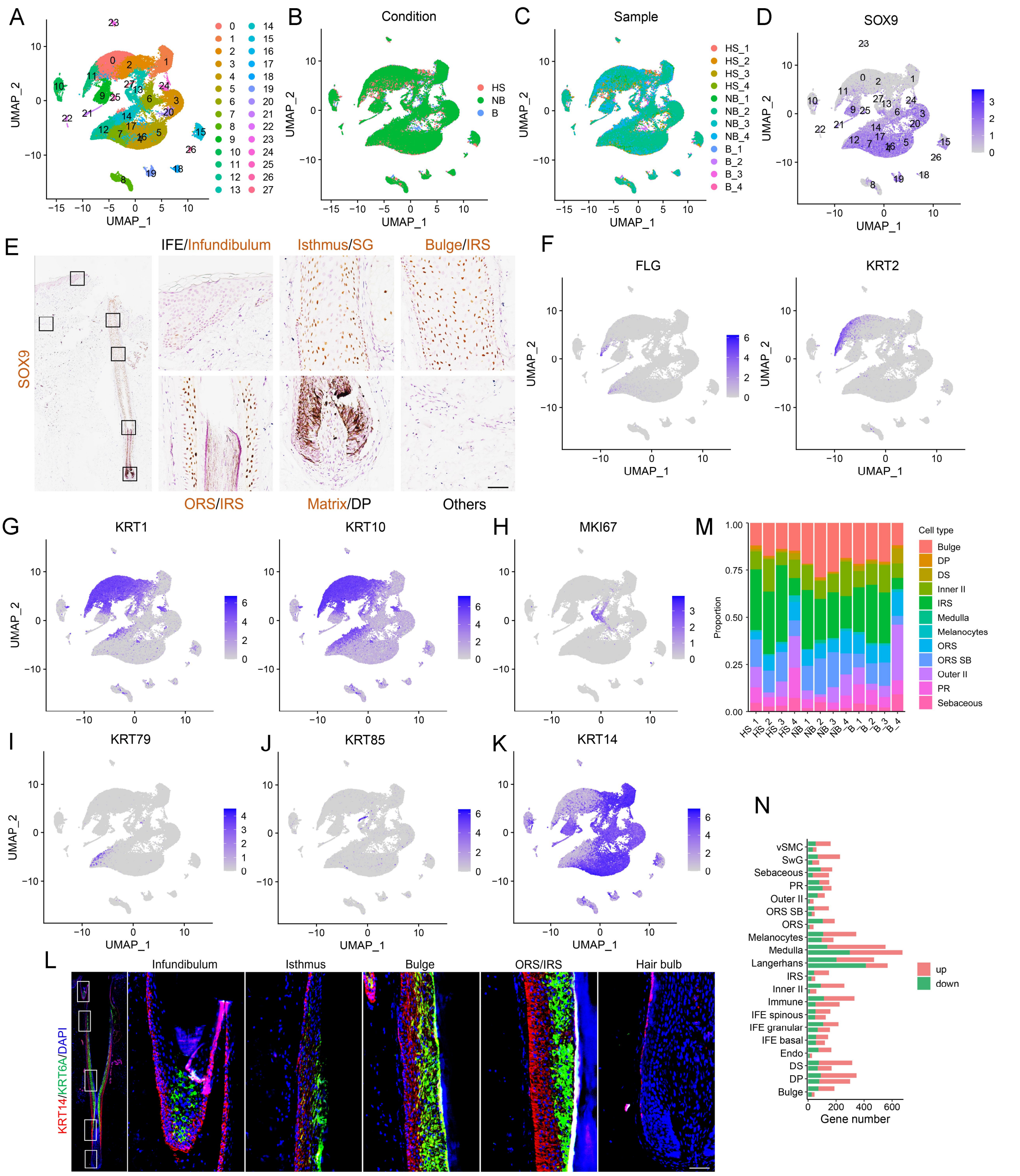


**Fig. S2: Cell type composition uncovered by scRNA-seq in HF units of AGA and healthy scalps.**

**A** Uniform manifold approximation and projection (UMAP) plot for all clusters. **B, C** Cells are colored by conditions and samples' identity in the UMAP plot. **D** Feature plot showing expression of the key hair follicle cell marker SOX9. **E** IHC staining of SOX9 on skin sections from healthy human scalp (n=3 individuals). **F-K** Feature plots showing the expression of the IFE granular (FLG, KRT2), IFE spinus (KRT1, KRT10), PR (MKI67), Inner II (KRT79), Medulla (KRT85), IFE basal (KRT14) markers. **L** Co-immunostaining of KRT14 and

KRT6A on skin sections from healthy human scalp (n=3 individuals). **M** Bar graph representing the proportion of major cell type populations of each sample. **N** Bar chart showing the differential gene number of major cell type populations. The upper bar of each cell type representing the differential gene number of balding compared with non-balding, and the other representing the differential gene number of balding compared with healthy HFs.

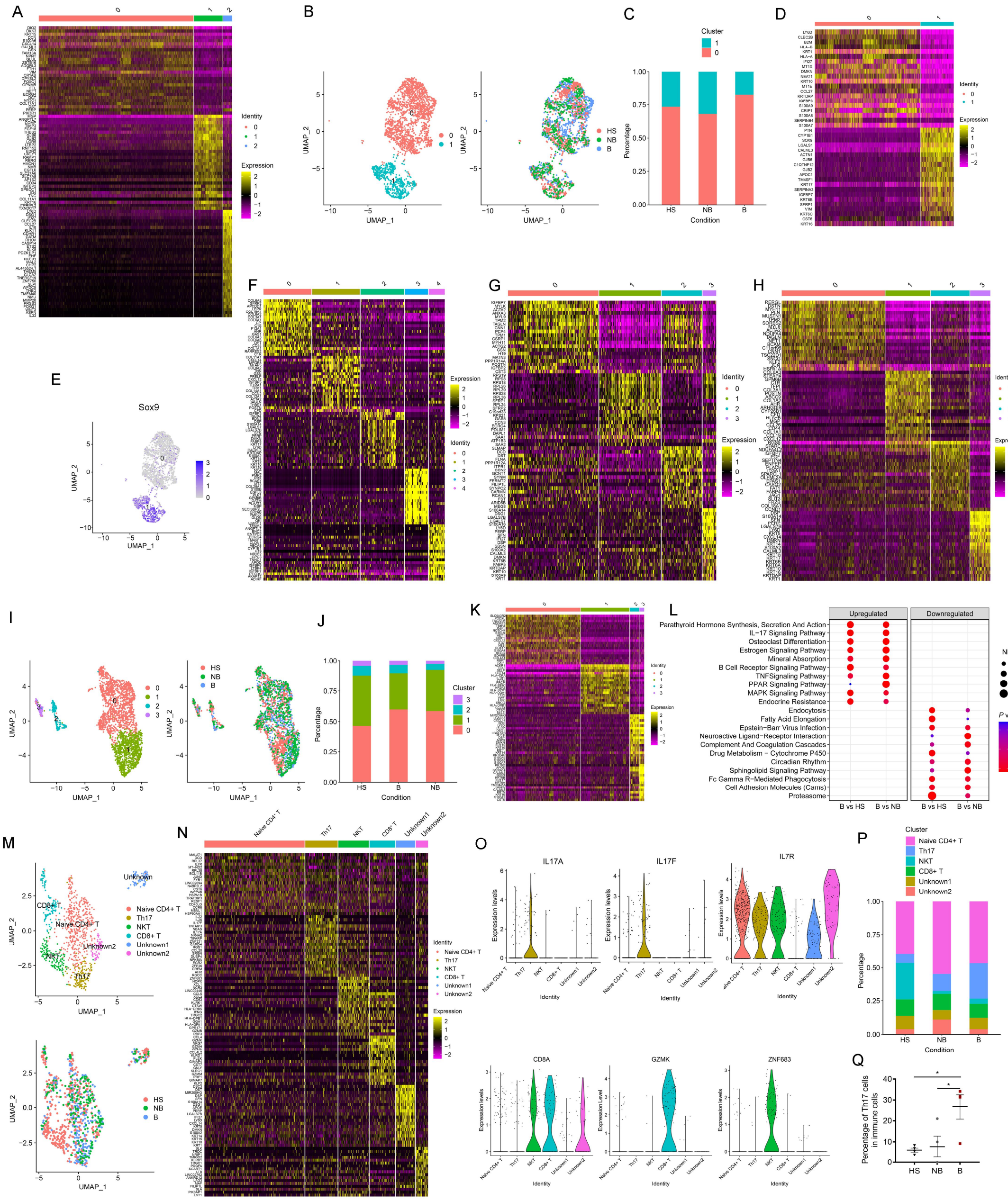


**Fig. S3: Identification of cell subpopulations in AGA and healthy scalps.** **A** Heatmap of top marker genes of bulge cells. **B** UMAP plot showing subclusters and sample conditions of PR cells. **C** Bar graph representing the percentage of subpopulations of PR cells. **D** Heatmap of top marker genes of PR cells. **E** Feature plot showing expression of the key hair follicle cell marker SOX9 in PR cells. **F-H** Heatmap of top marker genes of DP (F), DS (G) and vSMC (H) cells. **I** UMAP plot showing subclusters and sample conditions of endothelial cells. **J** Bar graph representing the percentage of subpopulations of endothelial cells. **K** Heatmap of top marker genes of endothelial cells. **L** Representative GO terms enriched in balding versus non-balding or healthy endothelial cells. NES stands for enrichment scores. The color keys from red to blue indicate the range of p value. **M** UMAP plots showing subclusters and sample conditions of immune cells. **N** Heatmap of top marker genes of immune cells. **O** Violin plots showing the expression of markers of Th17 markers (IL17A, IL17F), Naive $CD4^+$ T (IL7R), $CD8^+$ T (CD8A, GZMK) and NKT (ZNF683) cells. **P** Bar graph representing the percentage of different cell types of immune cells. **Q** The ratio of Th17 cells in HS, NB and B conditions. The data represent the means±SEM. *$p<0.05$, determined by unpaired (B vs HS) or paired (B vs NB) student's t-test.

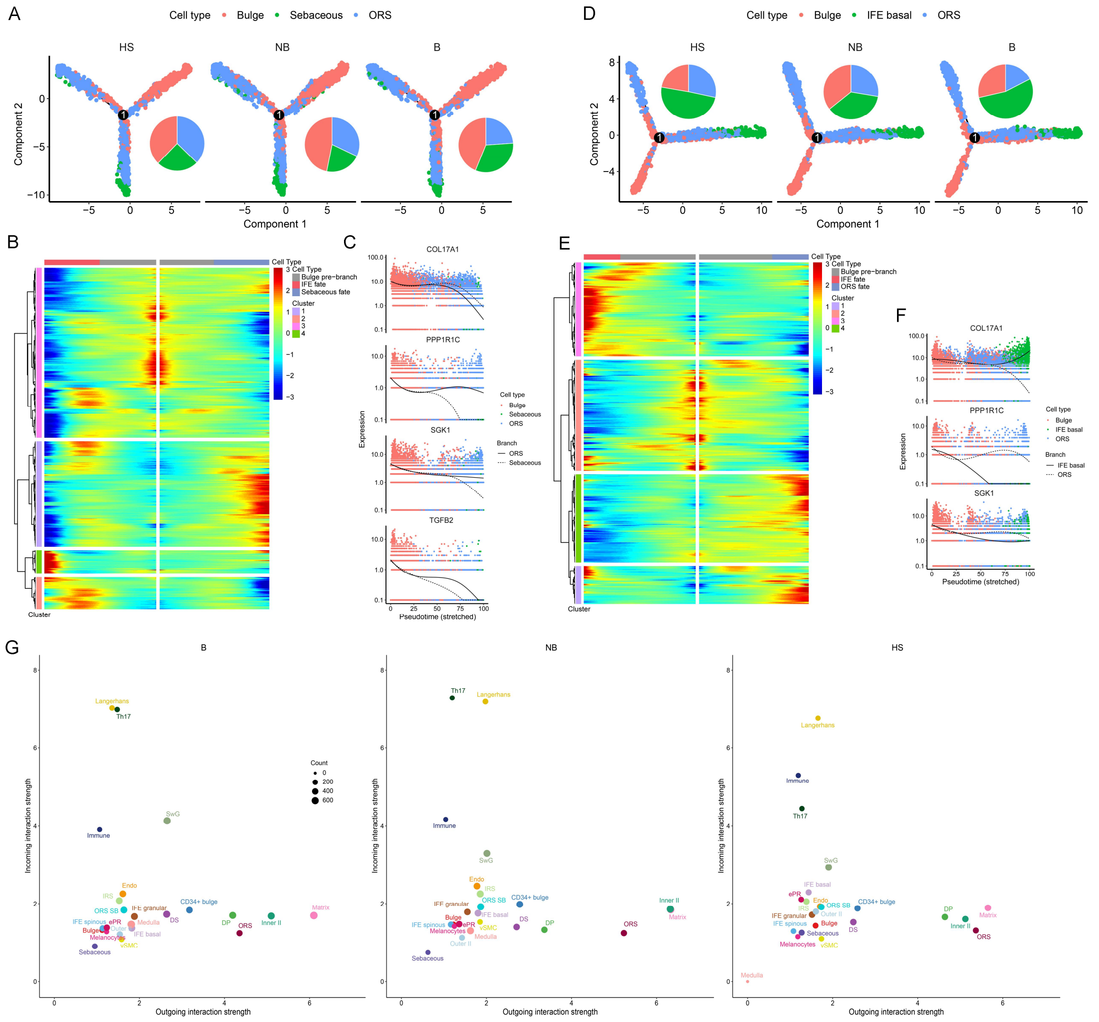


**Fig. S4: Changes in HFSC fate determination and cell-cell communications in AGA.**

**A** Pseudotemporal cell ordering of bulge, ORS and sebaceous gland cells along differentiation trajectories. Pie charts showing the cell ration on each differentiation branch. **B, C** Heatmap of top-scoring targets with differential expression across ORS fate branch and sebaceous fate branch (**B**) and the expression of critical differentiation markers (**C**) in pseudotime. **D** Pseudotemporal cell ordering of bulge, ORS and IFE basal cells along differentiation trajectories. Pie charts showing the cell ration on each differentiation branch. **E, F** Heatmap of top-scoring targets with differential expression across ORS fate branch (**E**) and IFE fate branch and the expression of critical differentiation markers (**F**) in pseudotime. **G** Scatter plots showing incoming/outgoing interaction strength for all HF cell types.

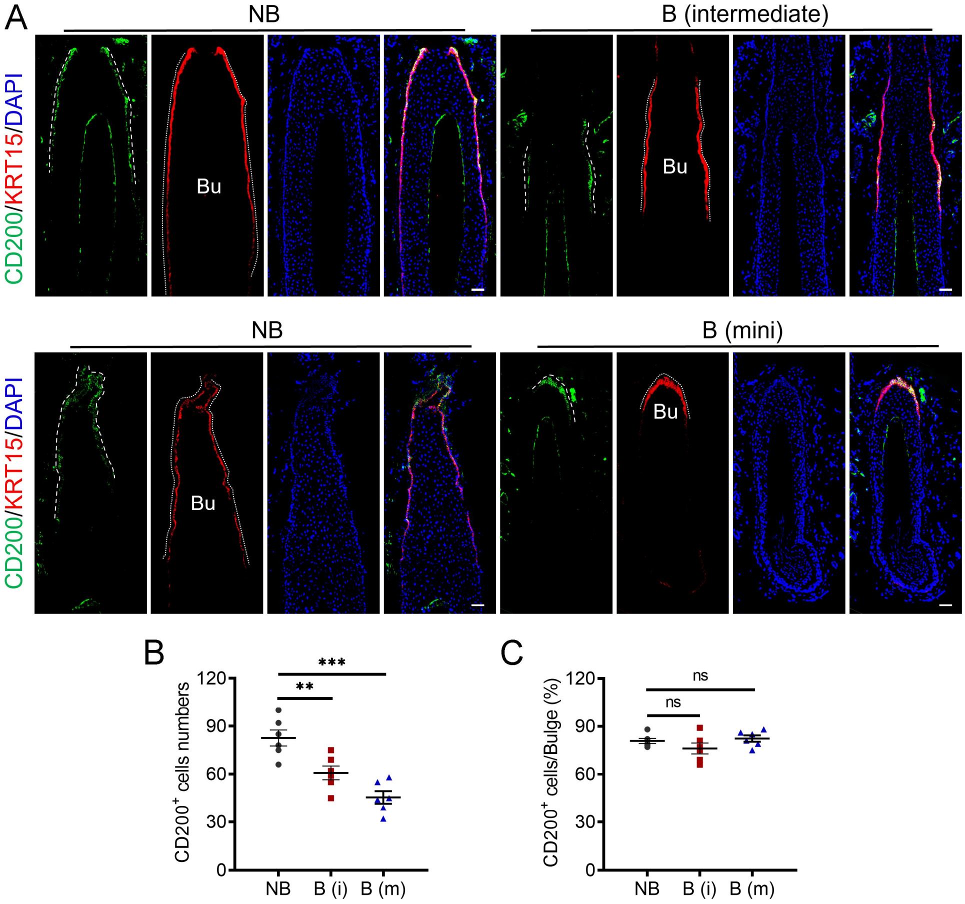


**Fig. S5: The percentage of HFSCs is not affected in AGA anagen HFs.**

**A** Representative immunostaining images of CD200 and K15 on terminal anagen HFs from non-balding occipital scalps and intermediate/mini anagen HFs from balding frontal scalps of AGA patients (n=6 anagen HFs from 3 patients). **B, C** Quantification of $CD200^{+}$ stem cells numbers (**B**) and their proportion (**C**) in bulge cells. Data are presented as mean±SEM and were analyzed by one-way ANOVA with Tukey's post hoc test. **p<0.01, ***p<0.001. ns, not significant.

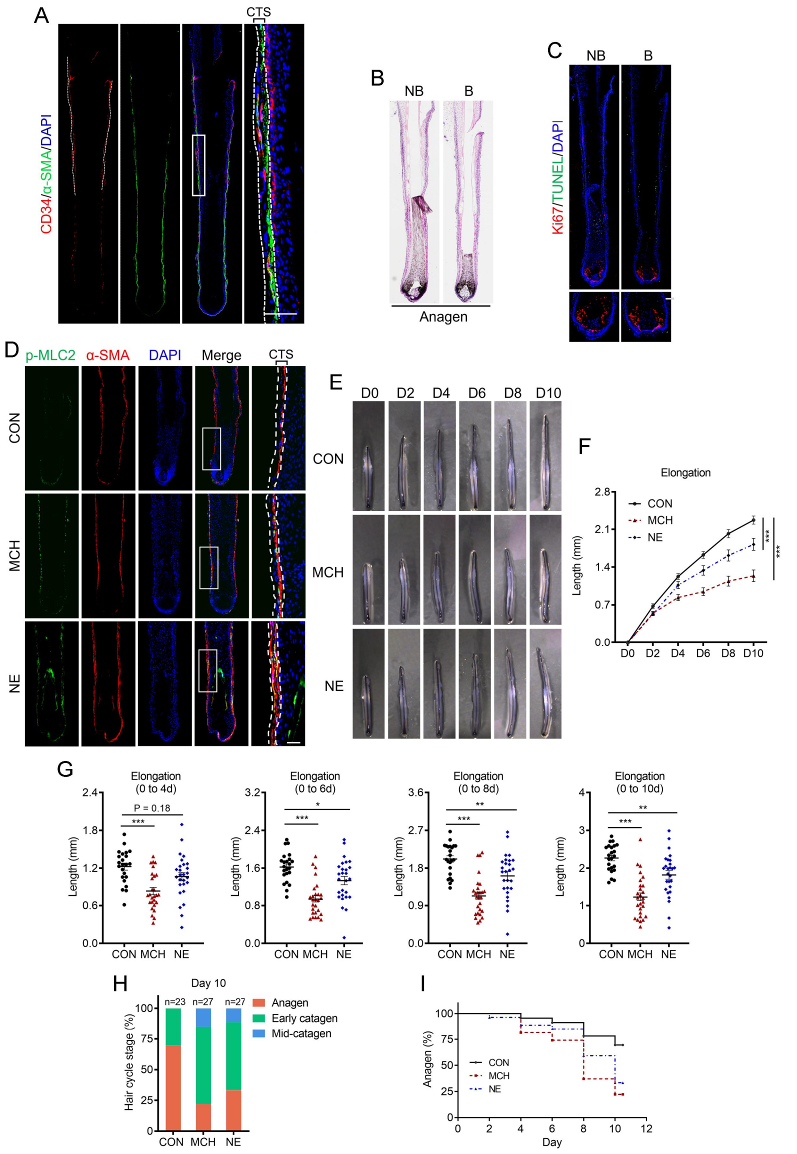


**Fig. S6: Aberrant activation of CTS contraction induces premature hair regression.**

**A** Immunostaining shows the location of CTS cells (DS and vSMC cells) and progenitor cells labeled by α-SMA and CD34 respectively (n=3 individuals). **B** Identification of anagen HFs in using HE staining. **C** Co-immunostaining of Ki67 and TUNEL with anagen HFs. **D** Co-immunostaining of p-MLC2 and α-SMA revealed the activation of CTS contraction in HFs treated with MCH (10 μM) and NE (0.1 μM) (n=6 HFs from 3 individuals). CTS is indicated by the two dotted lines. **E** Representative images of hair shaft elongation of human HFs ex vivo with MCH and NE as indicated (n=23/27/27 HFs; from 3 individuals). **F**

Quantification of hair shaft elongation on day 2/4/6/8/10 with MCH and NE treatments. **G** Scatter plots show the elongation length of hair shaft with MCH and NE treatments on day 4/6/8/10. **H** Macroscopic quantification of hair cycle stage of HFs with MCH and NE treatments on day 10. **I** Percentages of HFs in anagen with MCH and NE treatments on day 2/4/6/8/10. Data are presented as mean±SEM and were analyzed by one-way ANOVA with Tukey's post hoc test (G) and two-way ANOVA with a post hoc Holm–Sidak's multiple comparisons test (F). *p<0.05, **p<0.01, ***p<0.001.

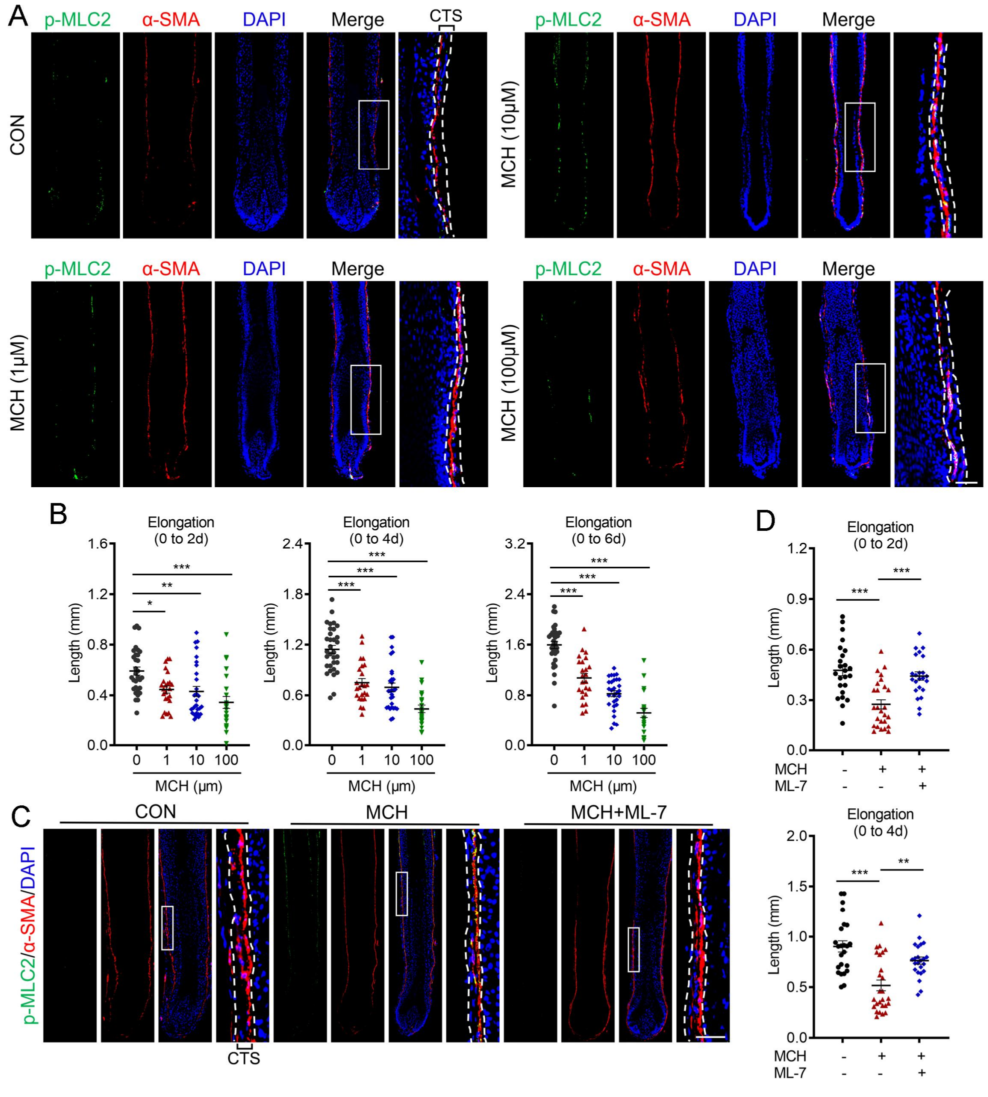


**Fig. S7: MCH-induced premature hair regression relies on CTS contraction.**

**A** Immunostaining of p-MLC2 and α-SMA revealed the activation of muscle contraction in anagen HFs treated with MCH at concentration of 1 μM, 10 μM, 100 μM as indicated (n=5 HFs in each group from two donors). **B** Scatter plots show the elongation length of hair shaft with MCH at different concentrations on day 2/4/6. **C** Representative immunostaining of p-MLC2 and α-SMA on HFs treated with MCH and ML-7 as indicated (n=6 HFs in each group from two donors). **D** Scatter plots show the elongation length of hair shaft with MCH and ML-7 as indicated on day 2. CTS is located between the two dotted lines. Data are presented as mean±SEM and were analyzed by one-way ANOVA with Tukey's post hoc test. *p<0.05, **p<0.01, ***p<0.001.

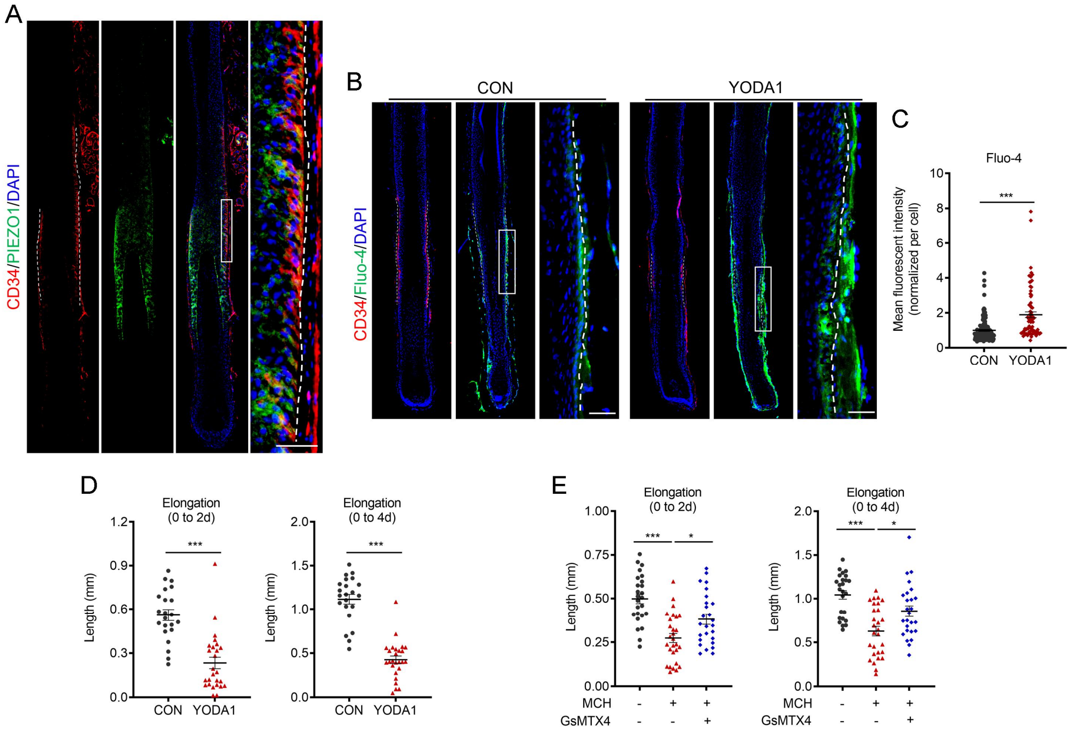


**Fig. S8: CTS contraction drives premature hair regression via PIEZO1 signaling.**

**A** Co-immunostaining of CD34 and PIEZO1 on anagen HFs from balding scalp of AGA patients (n=3 individuals). **B, C** (**B**) CD34 and Fluo-4 staining were performed on serial sections of HFs treated with or without YODA1 (n=3) and (**C**) the Fluo-4 mean fluorescent intensity in $CD34^{+}$ progenitor cells was quantified (n=80-100 cells from 3 different HFs). **D** Scatter plots show the elongation length of hair shaft with YODA1 treatment on day 2 and day 4. **E**

Scatter plots show the elongation length of hair shaft with MCH and GsMTX4 treatments on day 2 and day 4. Data are expressed as mean±SEM and were analyzed by two-tailed unpaired Student’s t-test. *p<0.05.***p<0.001.

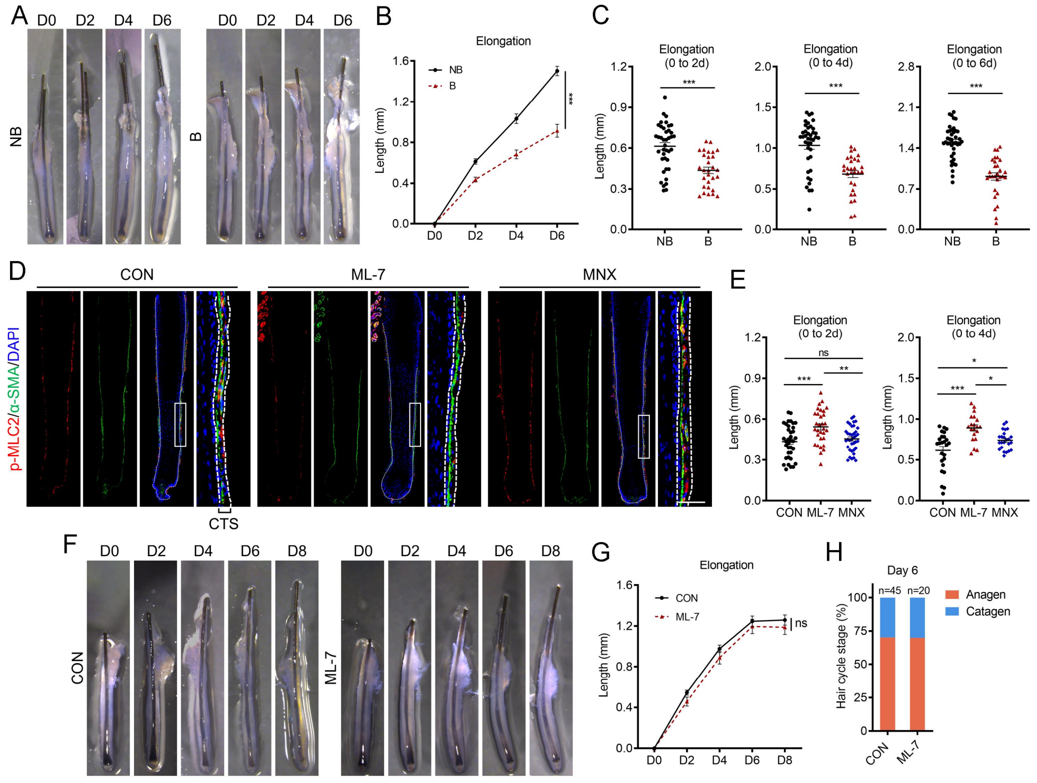


**Fig. S9: ML-7 shows a better effect on improving the growth of balding HFs from AGA patients.**

**A** Representative images of hair shaft elongation of anagen HFs from frontal balding (B) and occipital non-balding (NB) scalps of AGA patients in ex vivo culture system (n=29/40 HFs from 4 AGA patients for each group). **B** Quantification of hair shaft elongation on day 2/4/6. **C** Scatter plots show the elongation length of hair shaft on day 2/4/6. **D** Co-immunostaining of p-MLC2 and α-SMA revealed the activation of CTS contraction in balding HFs treated with ML-7 (0.3 μM) and MNX (100 μM) (n=6 balding HFs from 3 patients). Right panels, magnified images of boxed areas. Scale bar: 50 μm. **E** Scatter plots show the elongation length of hair shaft with ML-7 and MNX treatments on day 2/4. **F** Representative images of hair shaft elongation of NB HFs from AGA patients in ex vivo culture system treated without or with ML-7 (n=45/20 HFs from 4 AGA patients for each group). Of note, the HFs treated with and without ML-7 both exhibited typical early catagen morphology at day 6 to day 8. **G** Quantification of hair shaft elongation on day 2/4/6/8. **H** Macroscopic quantification of hair cycle stage of HFs with indicated treatments on day 6.Data are expressed as mean±SEM and were analyzed by one-way ANOVA with Tukey's post hoc test (E), two-way ANOVA with a post hoc Holm–Sidak's multiple comparisons test (B, G), and two-tailed unpaired Student's t-test (C). *$p<0.05$, **$p<0.01$, ***$p<0.001$. ns, not significant.
